# Cilia to basement membrane signalling is a biomechanical driver of autosomal dominant polycystic kidney disease

**DOI:** 10.1101/2024.06.06.597723

**Authors:** Manal Mazloum, Brice Lapin, Amandine Viau, Rushdi Alghamdi, Martine Burtin, Pascal Houillier, Lydie Cheval, Gilles Crambert, Amandine Aka, E. Wolfgang Kuehn, Camille Cohen, Stéphanie Descroix, Tilman Busch, Michael Köttgen, Serge Garbay, Marie-Christine Verpont, Brigitte Lelongt, Sylvie Coscoy, Fabiola Terzi, Frank Bienaimé

**Affiliations:** Université Paris Cité, INSERM U1151, CNRS UMR8235, Institut Necker Enfants Malades-INEM, Département « Croissance et Signalisation », Paris, France; Hôpital Universitaire Lapeyronie, service de Néphrologie Dialyse et Transplantation, Montpellier, France; Institut Curie, Université PSL, Sorbonne Université, CNRS UMR168, Laboratoire Physico-Chimie Curie, 75005 Paris, France; Université Paris Cité, Imagine Institute, Laboratory of Hereditary Kidney Diseases, INSERM UMR1163, Paris, France; Department of Clinical Physiology, Faculty of Medicine, King Abdulaziz University, Jeddah, Saudi Arabia; Sorbonne Université et Université Paris Cité, Centre de Recherche des Cordeliers, INSERM, Paris, France; Assistance Publique-Hôpitaux de Paris, Hôpital Européen Georges Pompidou, Service de Physiologie, Paris, France; Renal Division, Department of Medicine, Medical Center, Faculty of Medicine, University of Freiburg, Freiburg, Germany; CIBSS - Centre for Integrative Biological Signalling Studies, Freiburg, Germany; Université Paris Cité, Centre de recherche pour l’inflammation, INSERM U1149, EMR8252, Paris, France and service de Néphrologie, Hôpital Bichat, Assistance publique Hôpitaux de Paris, Paris, France; Sorbonne Université, CoRaKiD, INSERM UMRS1155, Hôpital Tenon, Paris, France; Assistance Publique-Hôpitaux de Paris, Hôpital Necker Enfants Malades, Service de Physiologie, Paris, France

**Keywords:** Cystogenesis, tubular basement membrane, primary cilia, biomechanics

## Abstract

Autosomal dominant polycystic kidney disease (ADPKD), which affects around 4 million patients worldwide, is characterized by the formation of multiple tubule derived cysts, which grossly enlarge both kidneys and progressively compromise renal function. ADPKD mainly results from mutations in *PKD1*, leading to the loss of polycystin-1 protein, which localizes to primary cilia. Primary cilia are required for cyst formation but the biomechanical changes underlying cystogenesis upon loss of polycytin-1 are unknown. We find that cilia and polycystin-1 shape the tubular basement membrane (TBM). Combining orthologous mouse models with a tubule-on-chip approach allowing manipulations of TBM stiffness, we find that cilia regulate the composition and biomechanical properties of the TBM. In the setting of polycytin-1 loss, reduced TBM stiffness and increased luminal pressure act as biomechanical drivers of cyst formation. These findings suggest a novel biomechanical model for ADPKD and unveil that cilia to TBM signalling controls kidney shape.

## INTRODUCTION

Autosomal dominant polycystic kidney disease (ADPKD) is the most common inherited renal disorder, accounting for 5 to 10% of kidney failure cases^1^. ADPKD is characterized by the transformation of a subset of renal tubules into cysts, which increase in number and size leading to renal mass growth of up to 20-fold. As a result, kidney function is progressively lost with the majority having experienced kidney failure by the sixth decade^1^. Modest therapeutic effects are gained from blood pressure control and the vasopressin receptor V2R antagonist tolvaptan, the only approved treatment for ADPKD^2^.

ADPKD is caused by mutations that inactivate a complex formed by polycystin-1 (PC1, encoded by *PKD1*), a large orphan receptor, and polycystin-2 (PC2, encoded by *PKD2*) a cation channel. The PC1/PC2 complex principally localizes to primary cilia, filiform cell organelles protruding from the apical surface of renal tubular cells that integrate chemical and mechanical cues to modulate cell behaviour. Cystogenesis in ADPKD occurs in a recessive fashion at the cellular level, where a tubule cell with a heterozygous germline mutation in *PKD1* or *PKD2* acquires somatic inactivation of the remaining allele^3^. Yet, the mechanisms for how cysts form have not been resolved.

Cell proliferation sustains cyst growth downstream of polycystin inactivation, but is insufficient on its own to prompt cyst formation. Indeed, tubular cell proliferation increases in response to nephron reduction, ischemia reperfusion injury or albuminuria without causing cyst formation^4–6^. Defects in planar cell polarity (PCP) signalling have been proposed to cause cyst formation through a loss of oriented cell division and/or extension convergence^7,8^. However, recent evidence indicates that PCP defects are not sufficient to precipitate cyst formation^9,10^. Besides, multiple signalling pathways have been shown to promote cyst growth (*e.g*., mTOR activation, cAMP signalling, metabolic rewiring), but none of these seem activated during early tubule dilation^11^. Thus, the primary morphogenetic events causing tubule dilation are still only partially understood.

The primary cilium is instrumental in the process of cystogenesis. Ciliary expression of the PC1/2 complex is required to prevent cyst formation^12^. Genetic cilia ablation prevents cyst formation in *Pkd1* mutant mice^11,13^, thereby supporting the existence of a “cilia-dependent cyst activation” (CDCA) pathway, the nature of which has remained unclear so far. Two cilia-regulated genes promoting cystogenesis have been identified: *Ccl2*^13^ and *Cdk1*^14^, which are involved in macrophage recruitment and tubular cell proliferation, respectively. However, unlike cilia ablation, their inactivation does not prevent early dilation of *Pkd1*-deficient tubules but rather slows cyst expansion^13,14^, suggesting the existence of additional ciliary functions in cystogenesis.

In higher metazoans, the extracellular matrix (ECM) secreted by cells is a critical determinant of tissue geometry and mechanics. In mammalian kidneys, decellularization experiments have shown that the ECM, which mainly consists in juxtaposed tubular basement membranes (TBM), retains not only the shape of nephrons but also tubule compliance, which accommodates to the luminal pressure required to maintain kidney function^15^. The TBM consists of interacting collagen IV and laminin networks, which are linked by heparan sulfate proteoglycans and nidogen^16^. Because of their long hydrophilic carbohydrate chains, peptidoglycans such as heparan sulfate proteoglycans (HS) promote basement membrane (BM) hydration, which increases deformability^17^. In contrast, the collagen and laminin networks provide rigidity, which is increased by covalent bridges between collagen IV chains^18,19^. The biomechanical properties of the BM are thus largely determined by its thickness, composition and degree of crosslinking^20^.

The role of TBM mechanics in ADPKD has not been systematically investigated. The fact that intratubular pressure was not raised in cysts of rodent PKD models led pioneer researchers to propose that cysts may arise from increased TBM deformability^21^. Yet, early studies investigating this hypothesis, mostly relying on non-orthologous models of the disease, failed to show consistent modifications of TBM mechanics, structure or composition^21,22^.

Here, we combined tubule-specific *Pkd1*-deletion mouse models, with or without concomitant cilia ablation, and a tubule-on-chip model with tunable ECM properties to decipher the interaction between cilia, *Pkd1*, TBM and tubule dilation.

## RESULTS

### *Pkd1* loss triggers cilia-dependent proximal and distal nephron dilation through distinct mechanisms

To gain insights into the early morphogenic events underlying tubule dilation, we studied mice with a post-developmental inactivation of *Pkd1* (*Pkd1*^ϕ.tub^), at a very early stage of the disease that is 2 weeks after the completion of doxycycline treatment (*i.e.*, 8 weeks of age), when tubules display only slight dilations (**Figure 1A**). Morphometric analysis confirmed that *Pkd1* loss induces an increase in cross-sectional areas of proximal (PT) and distal tubules [consisting of collecting ducts (CD) and distal convoluted tubules (DCT)], which was prevented by concomitant cilia disruption following *Kif3a* or *Ift20* inactivation (**Figure 1A-B; Extended data figure 1A-C**). *Pkd1* inactivation caused a more pronounced increase in tubule cross-sectional area in distal tubules than in PT (**Figure 1C**; **Extended data figure 1D**). KI67 labelling revealed that PC1 deficiency increased cell proliferation in PT (**Figure 1A, D**), but not in the distal nephron (**Figure 1A-D; Extended data figure 1A-E**). Instead, we observed increased tubular cell stretching in CD and DCT, as reflected by an increase in the mean distance between two adjacent nuclei (later referred as internuclear distance; **Figure 1E, Extended data figure 1F**). Linear regression analysis showed that distension (*i.e.,* increase in internuclear distance) was the main factor explaining distal nephron dilation at this stage (**Figure 1F**; **Extended data figure 1G-H**). At a later time point, distension and increased proliferation were observed in both proximal and distal *Pkd1*^-/-^ tubules (**Figure 1G-J**). Genetic cilia ablation (*i.e.*, *Kif3a* or *Ift20* inactivation) rescued PC1-deficient distal nephron distension, tubular cell proliferation and cyst development (**Figure 1G-J**). Collectively, these results identify cilia-driven tubule distension as an important factor responsible for the early dilation of PC1-deficient tubules.

**Figure 1.**
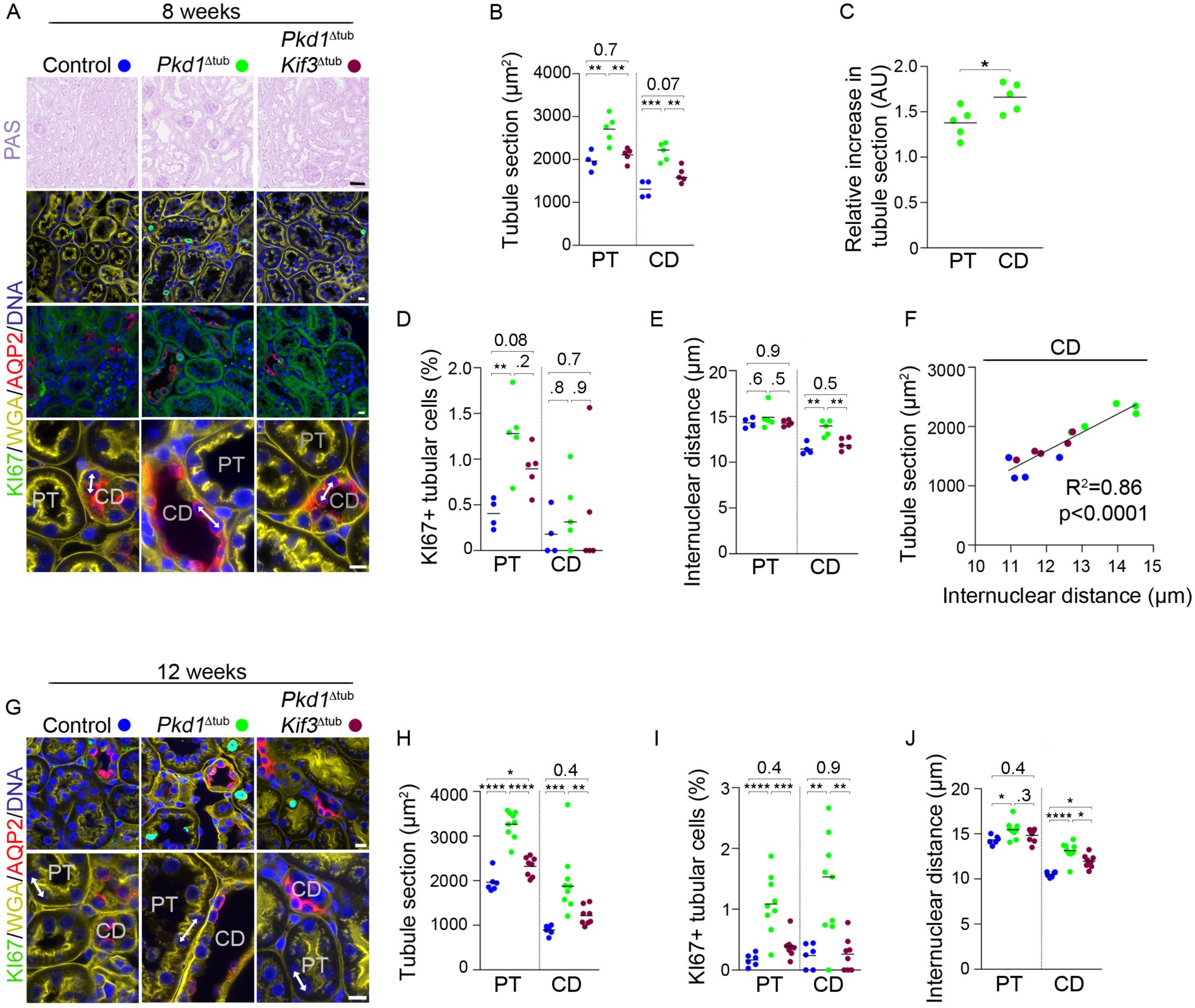
*Pkd1* deletion drives cilia-dependent distal tubule distension independently of cell proliferation. (**A**) Periodic acid Schiff’s (PAS) staining and labelling of KI67 (which stains proliferating cell), DNA, wheat germ agglutinin (WGA, which stains the brush border of proximal tubule [PT] and all basement membranes) and aquaporin 2 (AQP2, a collecting duct [CD] marker) of kidneys from 8-week-old control, *Pkd1*^Δtub^ and *Pkd1*^Δtub^; *Kif3a*^Δtub^ mice. Arrows: examples of internuclear distance measurements. Scale bars: 10 µm. (**B-E**) Quantification of mean PT and CD cross-sectional area (B), relative cross-sectional area increase in PT and CD from *Pkd1*^Δtub^ mice (*i.e.,* normalized to the mean of PT or CD cross-sectional area of control mice; C), proliferation index (percentage of KI67+ cells; D) and internuclear distance (E) in PT and CD of kidneys from 8-week-old control, *Pkd1*^Δtub^ and *Pkd1*^Δtub^; *Kif3a*^Δtub^ mice. (**F**) Linear regression of tubule cross-sectional area and internuclear distance for CD in the same mice groups at 8 weeks. (**G**) Labelling of KI67, DNA, WGA and AQP2 of kidneys from 12-week-old control, *Pkd1*^Δtub^ and *Pkd1*^Δtub^; *Kif3a*^Δtub^ mice. Arrows: examples of internuclear distance measurements. Scale bars: 10 µm. (**H-J**) Quantification of mean tubule cross-sectional area (H), proliferation index (KI67+ cells) (I) and internuclear distance (J) in PT and CD of kidneys from 12-week-old control, *Pkd1*^Δtub^ and *Pkd1*^Δtub^; *Kif3a*^Δtub^ mice. AU: arbitrary unit. Each dot represents one individual male mouse. One-way ANOVA followed by Tukey-Kramer test: *P<0.05, **P<0.01, ***P<0.001, ****P<0.01, or the indicated P-value.

### Preferential distal tubule distension correlates with specific changes of the TBM

As tubule shape is determined by TBM geometry^15^, we reasoned that CD distension could be linked to specific TBM properties. TBM stiffness depends on its thickness and composition: laminin and cross-linked collagen IV provide rigidity, while the sugar moieties of heparan- and chondroitin-sulfate (HS/CS) confer flexibility by increasing BM hydration^20^. Studying these parameters in basal conditions, we noticed that, compared to PT, CD had thinner BM with higher HS content, suggesting a higher compliance (**Figure 2A-B**), as historically suggested by pressure-volume measurements on isolated rabbit tubules^15^. We further observed that distal nephron distension in *Pkd1*^Δtub^ mice was associated with prominent TBM thinning (**Figure 2C**; **Extended data figure 2A**). TBM thinning also occurred in PT but to a smaller extent than in CD or DCT (**Figure 2C**). A careful inspection of multiple tubule sections in transmission electron microscopy failed to detect any perforation or tubular cell protrusion through the TBM. Cilia ablation abolished the ability of PC1-deficient tubular cells to induce TBM thinning (**Figure 2C**; **Extended data figure 2A**). We further observed a specific increase in HS immunoreactivity of the TBM of PC1-deficient tubules undergoing distension, which was prevented by cilia disruption, without notable changes in collagen IV labelling (**Figure 2D-E**; **Extended data figure 2B-C**). These results indicate that *Pkd1* loss is associated with a cilia-dependent remodelling of the TBM. They also suggest that intrinsic properties of distal nephron TBM may render this segment permissive to distension. Detailed transmission electron microscopy images of TBM are presented in **Extended data figure 3**.

**Figure 2.**
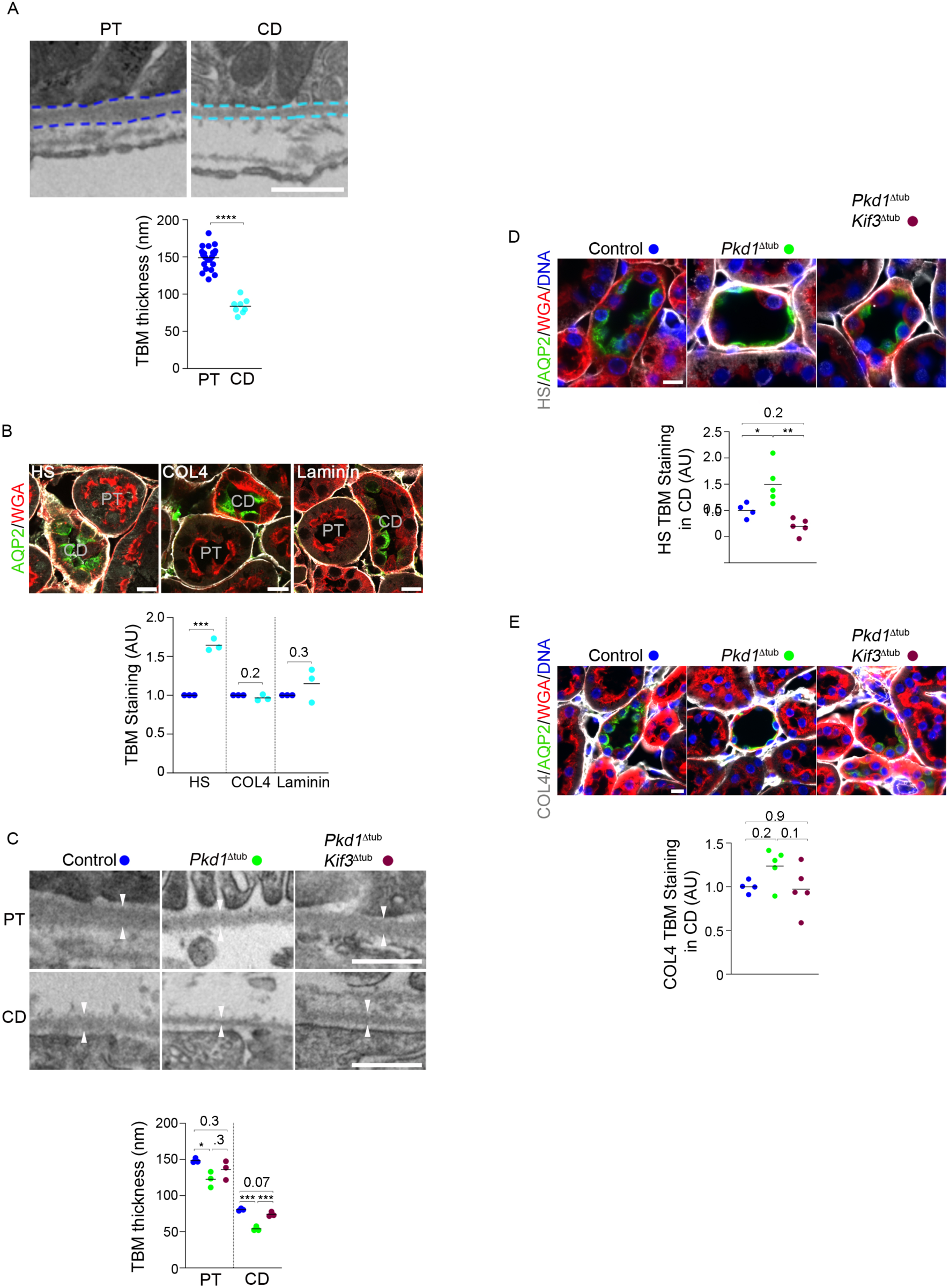
*Pkd1* deletion induces cilia-dependent basement membrane remodelling. (**A**) Transmission electron microscopy and quantification of proximal tubule (PT) and collecting duct (CD) tubular basement membrane (TBM) thickness in 8-week-old control mice. Dashed lines underline TBM. Each dot represents one individual tubule (mean of 3 to 21 measurements per tubule; n=3 mice). Scale bar: 0.5 µm. (**B**) Labelling of PT brush border (WGA), collecting duct (AQP2) and heparan sulfate (HS; left panel), collagen IV (COL4; middle panel) or laminin (right panel), and quantification of HS, COL4 and laminin signal intensity in 8-week-old control mice. Each dot represents one individual male mouse. Scale bar: 10 µm. Student t-test: ***P<0.001. (**C**) Transmission electron microscopy of TBM of PT and CD from 8-week-old control, *Pkd1*^Δtub^ and *Pkd1*^Δtub^; *Kif3a*^Δtub^ mice. Each dot represents one individual male mouse. Scale bar: 0.5 µm. (**D-E**) Staining and quantification of HS (D) and COL4 (E) in CD of the same animals. Each dot represents one individual male mouse. Scale bars: 10 µm. One-way ANOVA followed by Tukey-Kramer test: *P<0.05, **P<0.01, *** P<0.001 or the indicated P-value. AU: arbitrary unit.

### TBM remodelling and CD distension coincide with a specific transcriptional program

Our results suggest that cilia-dependent TBM remodelling promotes cyst formation in ADPKD and is decoupled from cell proliferation. Three non-mutually exclusive mechanisms can cause TBM distension without cell proliferation: (1) ECM digestion by proteases, which allows rapid BM remodelling during embryogenesis^23^; (2) incorporation of BM softeners such as hydrating HS/CS, which allows rapid BM remodelling during *C.elegans* development^24^; (3) deformation of the BM by traction forces exerted by cells through cytoskeleton motor proteins, as documented during cancer invasion^25,26^. To gain insights into the effectors involved, we profiled the transcriptome of PT and CD micro-dissected from 8-week-old *Pkd1*^Δtub^ and control mice (**Figure 3A-C**, **Extended data figure 4A**). The results are provided as **Supplementary file 1**. PT and CD specific gene expression profiles confirmed dissection quality (**Extended data figure 4B**). *Pdgfrb* and *Itgam* expressions were undetectable in CD, indicating the absence of contamination of the extracts by fibroblasts or macrophages, respectively. Overall, *Pkd1* inactivation induced a significant dysregulation in the expression of 1,409 genes in CD but only 357 in PT (**Extended data figure 4C**). Enrichment analyses showed an enrichment in gene sets related to cell proliferation in PC1-deficient PT, but not in CD; a finding consistent with our immunolabelling experiments (**Figure 3B**). More importantly, we identified a panel of 26 genes, which are upregulated in PC1-deficient CD but not in PT and are involved in the three aspects of BM remodelling that we prespecified (**Figure 3C**). These include 4 genes encoding ECM proteases, 3 genes allowing the production of BM softeners (*i.e.*, enzymes involved in HS/CS biogenesis; *Hs3st3al, Chsy1),* 6 additional ECM components and 13 genes involved in cell-BM interactions or cytoskeleton dynamics.

**Figure 3.**
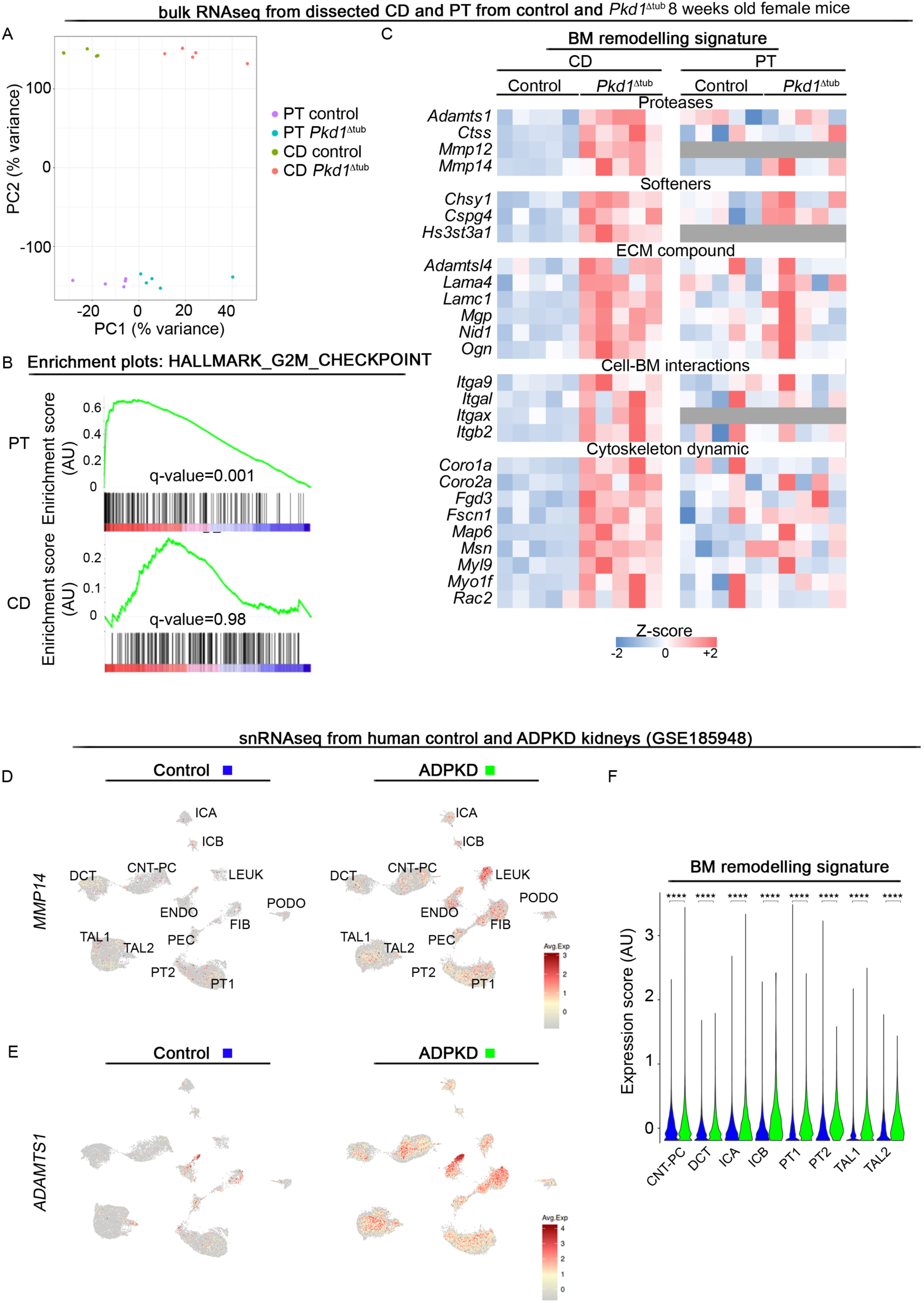
Transcriptomic profiling of micro-dissected proximal tubules (PT) and collecting ducts (CD) identifies a basement membrane (BM) remodelling signature associated with tubule distension. (**A**) Principal component analysis (PCA) of the transcriptome of proximal tubules (PT) and collecting ducts (CD) micro-dissected from control and *Pkd1*^Δtub^ 8-week-old female mice (n=5 mice per genotype, each dot represents a tubule segment from an individual male mouse). (**B**) Gene set enrichment analysis (GSEA) comparing proliferation related gene sets between control and mutant mice in PT or CD. (**C**) Heatmaps showing the relative expression for TBM remodelling candidate genes in CD and PT micro-dissected from control and *Pkd1*^Δtub^ 8-week-old female mice (n=5 mice per genotype). Grey in the heatmap of PT indicates undetectable expression. (**D-E**) Expression of *MMP14* (D) and *ADAMTS1* (E) in snRNAseq data of control and ADPKD human kidneys. PT proximal tubule, PEC parietal epithelial cells, TAL thick ascending limb of Henle’s loop, DCT distal convoluted tubule, CNT_PC connecting tubule and principal cells, ICA Type A intercalated cells, ICB Type B intercalated cells, PODO podocytes, ENDO endothelial cells, FIB fibroblasts, LEUK leukocytes, URO urothelium. (**F**) BM remodelling signature expression score in the indicated tubular cell population from control or ADPKD human kidneys. Bonferroni test : ****P<0.0001. AU: arbitrary unit.

Re-analysis of snRNAseq datasets comparing human ADPKD and healthy kidneys^27^ showed that the BM remodelling signature that we identified in distending CD from *Pkd1*^Δtub^ mice was significantly enriched in ADPKD human tubular cells (**Figure 3D-F**). Of note, in these datasets of terminal human ADPKD kidneys, TBM remodelling genes induction was not restricted to distal segments, suggesting a generalization of the process at late stages.

### Loss of *Pkd1* is associated with altered tubule mechanics

We were interested if remodelling of the TBM alters the physical properties of tubules. We isolated CD and PT from 8-week-old *Pkd1*^Δtub^ mice (with intact TBM) and measured their diameter variations in response to increasing luminal pressure using a dedicated isolated tubule perfusion setup (**Figure 4A**)^15^. We observed that PC1-deficient CDs, which have undergone distension but not proliferation (*i.e.*, dissected from 8-week-old mice; **Figure 1A-E**), showed a steep increase in tubule diameter at low pressure (*i.e.*, from 0 to 10 cmH_2_O; **Figure 4B-D**). In contrast, PC1-deficient PT, which have not yet undergone distension at this time point but are nonetheless larger than in control mice because of cell proliferation (**Figure 1A-E)**, showed similar pressure-diameter curves than control PT (**Figure 4E-F**). Repeating these experiments with PT isolated from 12-week-old animals, a time point where distension has occurred in this segment (**Figure 1G-J)**, we found a similar steep increase in tubule diameter than in distended PC1-deficient CD, although for a higher pressure threshold (*i.e.*, from 21 to 31 cmH_2_O; **Figure 4G-I**). Notably, decreasing pressure in distended PC1-deficient tubules resulted in a decrease of their diameter indicating elastic deformation **(Extended data figure 5**). Together, these results demonstrate that tubule distension and TBM remodelling in ADPKD are associated with an alteration of tubule mechanics, characterized by a steep increase in tubule diameter with segment-specific pressure thresholds.

**Figure 4.**
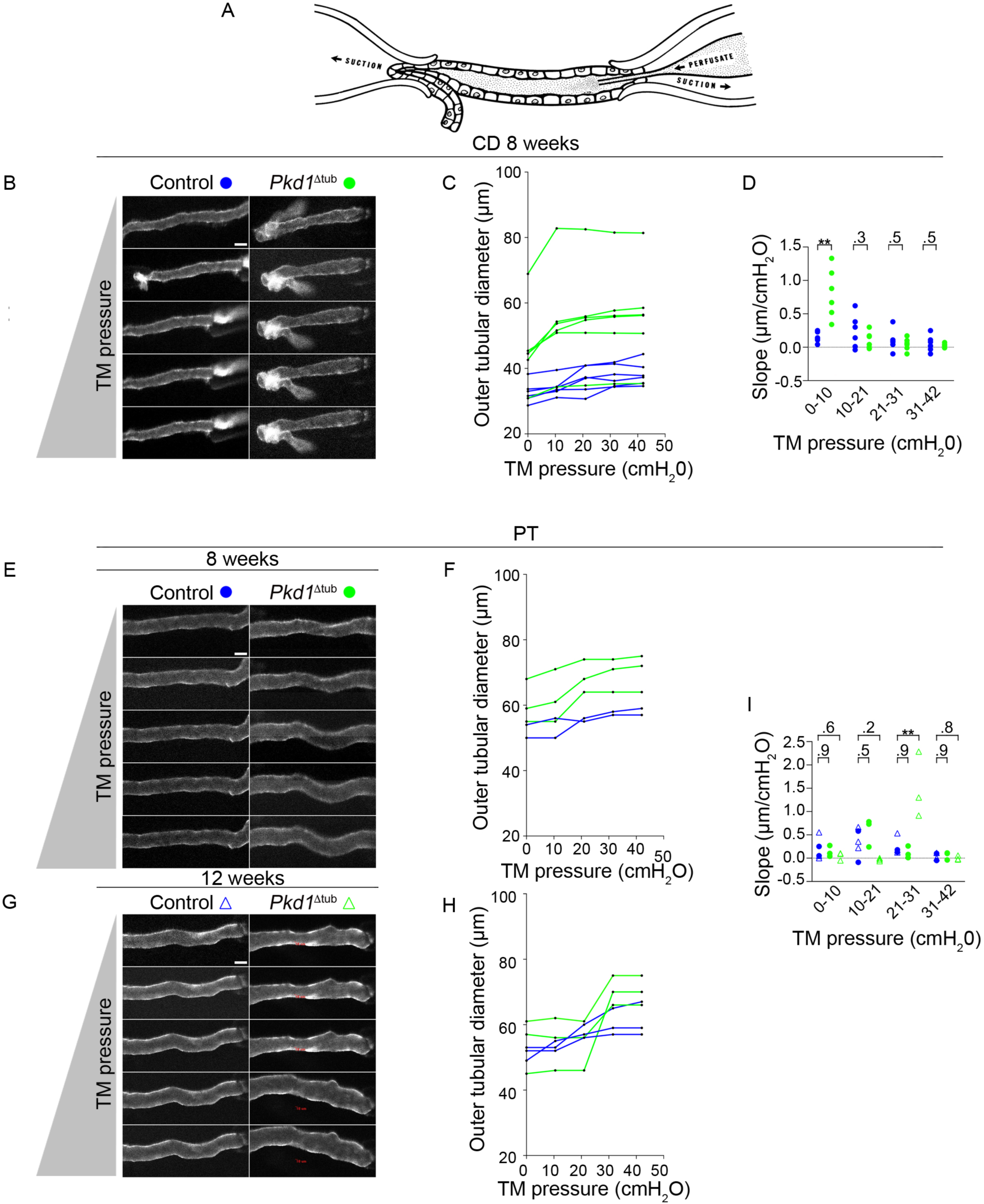
*Pkd1* deletion alters tubule biomechanics. (**A**) Scheme of isolated intact tubule perfusion setting^15^. (**B-C**) Representative images (B) and quantification of the variation in outer tubule diameter induced by a progressive increase in transmural pressure (TM; C) in collecting ducts isolated from 8-week-old control and *Pkd1*^Δtub^ male mice. Each dot represents one individual tubule (mean of 10 measurements per tubule) for a pressure increment. (**D**) Quantification of the slope of the diameter-pressure curves for each pressure increment in collecting ducts isolated from the same groups of mice. Each dot represents one individual tubule. (**E-F**) Representative images (E) and quantification of the variation in outer tubule diameter induced by incremental increase in TM pressure (F) in proximal tubules isolated from 8-week-old control and *Pkd1*^Δtub^ mice. Each dot represents one individual tubule (mean of 10 measurements per tubule) for a pressure increment. (**G-H**) Representative images (G) and quantification of the variation in outer tubule diameter induced by incremental increase in TM pressure (H) in proximal tubules isolated from 12-week-old control and *Pkd1*^Δtub^ mice. Each triangle represents one individual tubule (mean of 10 measurements per tubule) for a pressure increment. (**I**) Quantification of the slope of the diameter-pressure curves for each pressure increment in PT isolated from 8 and 12-week-old control and *Pkd1*^Δtub^ mice. PT from control animals at 8 (blue dot) and 12 weeks (blue triangle) were pooled in analyses. Each symbol represents one individual tubule. Scale bars: 50 µm. Student t-test: **P<0.01.

### Changes in TBM mechanical properties result in permissive conditions for cystogenesis

Our results regarding the early modification of TBM in ADPKD suggest that increased ECM deformability favours cystogenesis. This model contradict the general idea that increased ECM stiffness promotes tubular cell proliferation and kidney disease progression^28,29^. Yet, experimental manipulation of ECM stiffness has not been performed in ADPKD. To investigate this question *in vitro*, we used an established tubule-on-chip model, whereby immortalized tubular cells form ciliated tubules of physiological dimensions in a deformable ECM scaffold^30^. Using edited ciliated clonal lines derived from mouse collecting duct (mIMCD3)^31^, we observed only minor differences between control and *Pkd1*^-/-^ clones in the ability to dilate ECM scaffolds with a collagen content of 9.5 g/l (**Figure 5A**). However, by lowering collagen content, thereby reducing stiffness, we observed that *Pkd1*^-/-^ clones dilated more rapidly the ECM scaffold than the parental line (**Figure 5B-C**). This increased dilation was not caused by increased cell proliferation but by an increase in internuclear distance (**Figure 5D-F**). These *in vitro* results demonstrate that ECM mechanics affect the ability of *Pkd1*^-/-^ cells to deform tubules, so that ECM deformability facilitates dilation. To test this concept *in vivo*, we inactivated *Pxdn*, which encodes peroxidasin, a collagen IV specific cross-linking enzyme (which germinal ablation induces a 20% decrease in TBM elastic module)^32^, in *Pkd1*^Δtub^ mice (**Figure 5G-H**). While *Pxdn*^-/-^ mice have no kidney phenotype at baseline^32^, the modest alteration of TBM mechanics caused by *Pxdn* inactivation translated into increased cyst burden in *Pkd1*^Δtub^ mice (**Figure 5G-H**). At the studied time point, cystogenesis was more pronounced in males compared to females. Yet, the impact of *Pxdn* inactivation on cyst area was not affected by gender. These results are consistent with a model where reduced TBM stiffness facilitates tubule deformation and cystogenesis.

**Figure 5.**
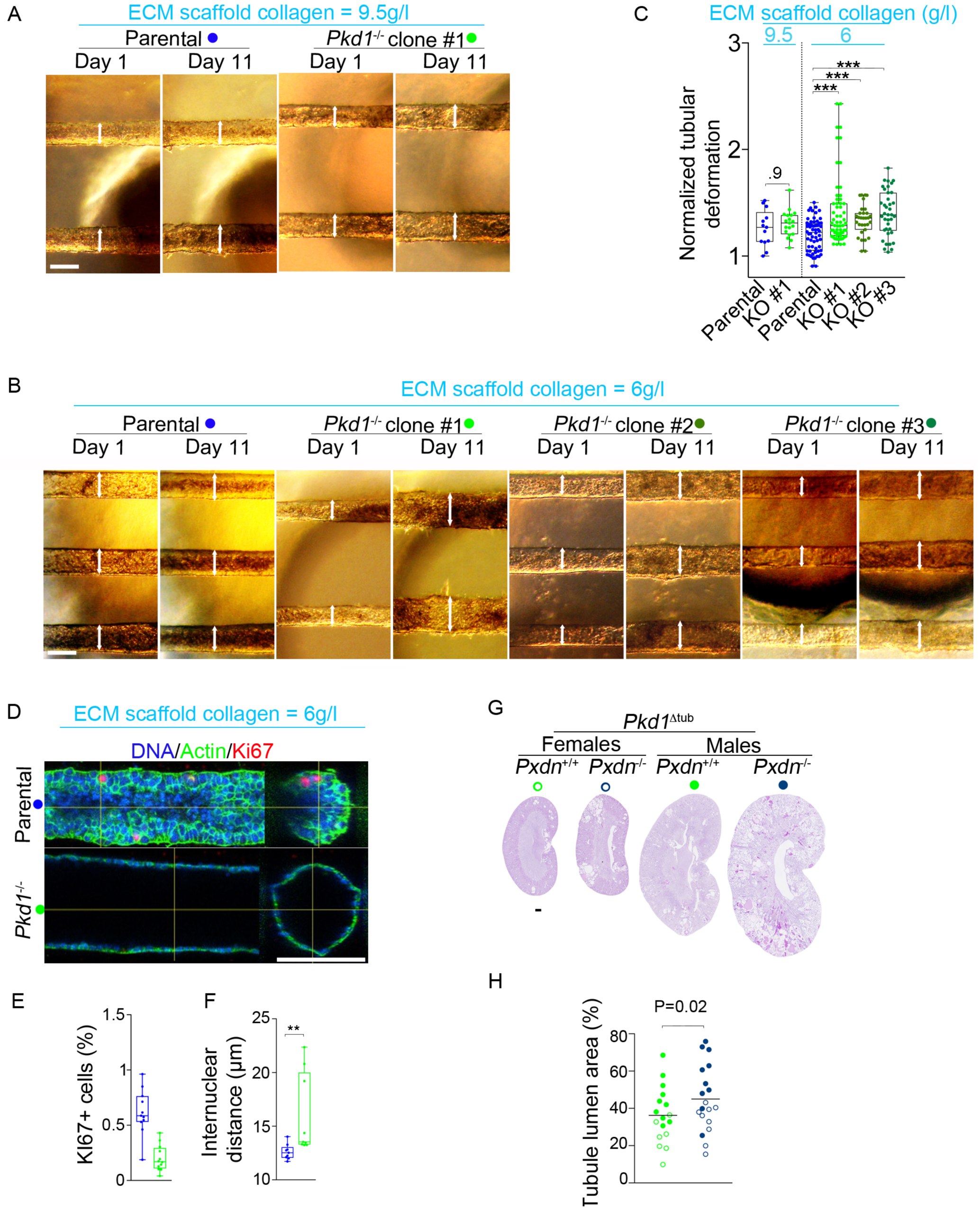
Extracellular matrix mechanics modulates *Pkd1*^-/-^ tubule-on-chip dilation *in vitro* and cyst formation *in vivo*. (**A-C**) Representative images (A-B) of tubules-on-chip formed by parental or *Pkd1*^-/-^ mIMCD-3 clones in scaffolds containing 9.5 (A) or 6 g/l (B) collagen I, 1 and 11 days after confluency and quantification (C) of the normalized tubule deformation at 11 days. Arrows indicate tubule diameter. Scale bar: 100 µm. (**D-F**) Labelling (D) and quantification of KI67 (E) and internuclear distance (F) in parental and *Pkd1*^-/-^ tubules-on-chip. Each dot represents one individual tubule. Mann-Whitney test: **P<0.01, ***P<0.001. (**G-H**) Periodic acid Schiff’s staining (G) and quantification (H) of tubular dilations of kidneys from 12-week-old *Pkd1*^Δtub^ female (open circle) and male (closed circle) mice with or without concomitant peroxidasin gene (*Pxdn)* inactivation. Each dot represents one individual mouse. Scale bar: 0.5 mm. P indicates the P-value for *Pxdn* genotype effect in Two-way ANOVA (gender and *Pxdn* genotype; P*-*value for interaction between gender and genotype = 0.8).

### Tubule obstruction exacerbates PC1-deficient tubule distension and precipitates cystogenesis

Tubular obstruction from enlarging cysts has been proposed to contribute to new cyst formation and micro-puncture as well as microdissection studies have shown that cysts frequently arise from obstructed tubules^33^. We tested *in vivo* how alterations in TBM biomechanics affect tubule response to obstruction with respect to PC1 and cilia. After a single day of unilateral ureteral obstruction (UUO), *Pkd1*^Δtub^ mice showed a disproportional increase in kidney weight compared to controls. This was not the case in kidneys with concomitant cilia ablation (**Figure 6A-B**; **Extended data figure 6A**). The kidney weight increase correlated with enlargement of the CD and DCT, but not PT and was associated with a marked increase in internuclear distance, indicating tubule distension (**Figure 6C-H**; **Extended data figure 6B-D**). BrdU labelling excluded increased cell proliferation as a cause for CD dilation (**Extended data figure 6E-F**). Two-way ANOVA confirmed that a significant interaction between genotype and obstruction governs distal nephron distension **(Figure 6D-E**; **Extended data figure 6C-D)**.

**Figure 6.**
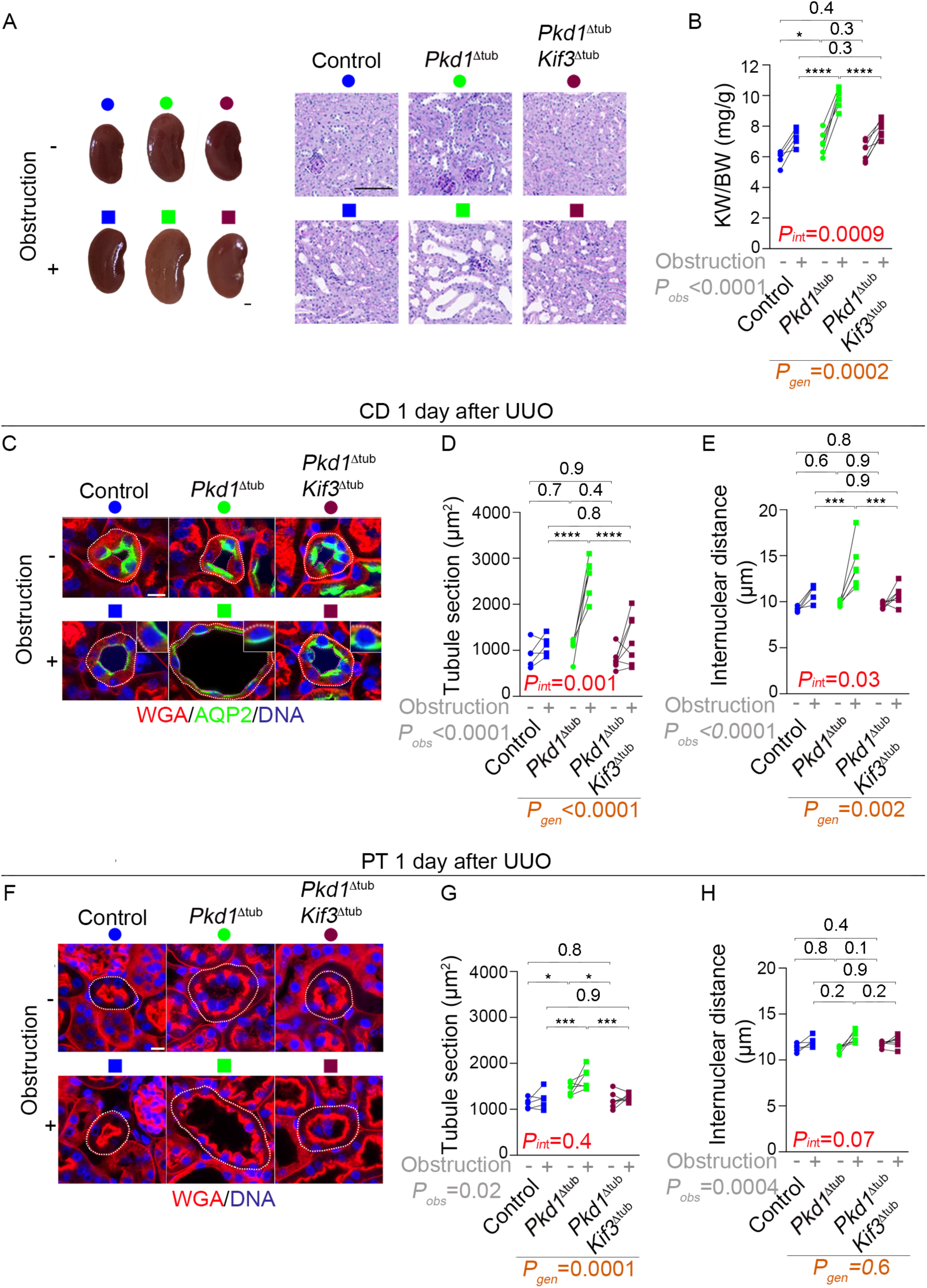
Ureteral obstruction accelerates *Pkd1*^Δtub^ distal tubule dilation in a cilia-dependent manner. (**A-B**) Kidney pictures, Periodic-acid Schiff’s (PAS) staining (A) and quantification (B) of kidney weight to body weight ratio (KW/BW) of obstructed and non-obstructed kidneys from 8-week-old control, *Pkd1*^Δtub^ and *Pkd1*^Δtub^; *Kif3a*^Δtub^ one day after unilateral ureteral obstruction (UUO). Scale bars: 1 mm (left) and 0.1 mm (right). (**C-H**) labelling of DNA, wheat germ agglutinin (WGA, which stains the brush border of proximal tubule [PT] and all basement membranes) and aquaporin 2 (AQP2, a collecting duct [CD] marker; C, F), quantification of mean tubule cross-sectional area (D, F) and internuclear distance (E, H) of CD (C-E) or PT (F-H) of kidneys from the same animals. Each pair of linked symbols (dot and square) represents the obstructed (square) and non-obstructed (dot) kidneys of an individual female mouse. Scale bars: 10 µm. Two-way ANOVA, P-value for obstruction (grey; *P_obs_*), genotype (brown; *P_gen_*) and their interaction (red; *P_int_*), followed by Tukey-Kramer test: *P<0.05, **P<0.01, ***P<0.001, ****P<0.0001.

Prolonging obstruction resulted in cystic transformation of PC1-deficient kidneys, with a two- and three-fold increase in kidney weight to body weight ratio 4 and 14 days after the surgery, respectively (**Figure 7A-D; Extended data figure 7A-B**). Cilia ablation reduced obstructed PC1-deficient kidneys enlargement to the mild level observed in cilia-deficient mice with intact *Pkd1* (*i.e.*, *Kif3a*^Δtub^; **Figure 7C-D**). Cystic transformation of *Pkd1*-deficient kidneys was mainly the consequence of an increase in DCT and CD cross-sectional area, while tubule internuclear distance remained higher than in control or PC1-deficient DCT and CD lacking cilia (**Figure 8A-C; Extended data figure 7C-H**). We then asked if the cystic transformation of PC1-deficient tubule could be caused by an excessive proliferative response to obstruction. As expected, labelling cycling cells with PCNA or KI67 revealed that, with time, obstruction increased the fraction of cycling cells in CD and DCT. However, the magnitude of this increase was similar across genotypes **(Figure 8D, Extended data figure 8A-D**). Consistent with our baseline findings, non-obstructed PT lacking PC1 displayed a higher fraction of cycling cells than control or deciliated PT. Obstruction further increased the cycling cell fraction in all genotypes (**Extended data figure 8C-E**). The proliferation rate did not correlate with PT deformation: obstruction reduced PT cross-sectional area in control animals, indicating atrophy. This compaction of PT was similarly prevented by *Pkd1* or cilia ablation (**Figure 8G-H)**.

**Figure 7.**
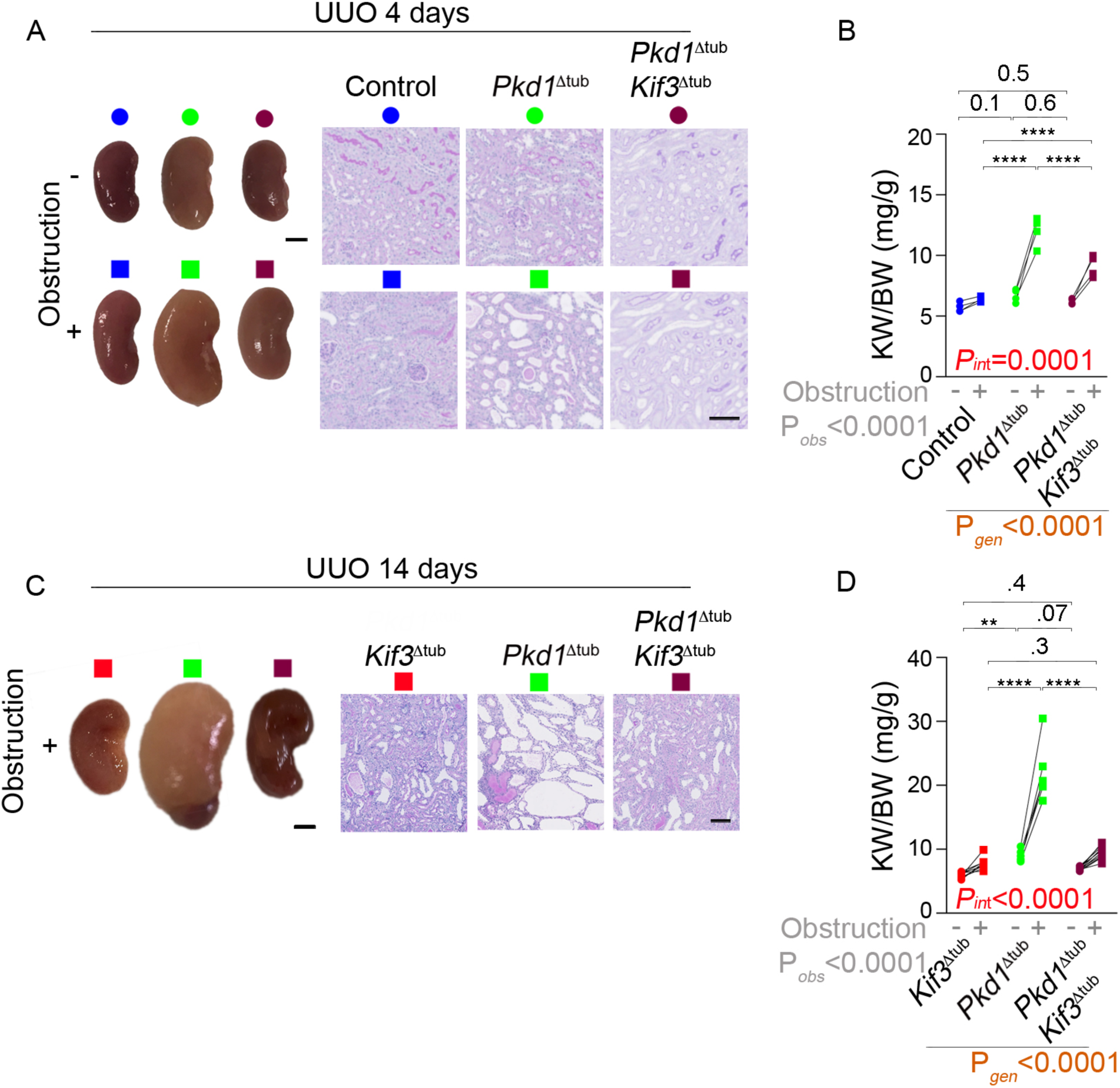
Ureteral obstruction triggers explosive cystogenesis in *Pkd1*^Δtub^ distal tubules in a cilia-dependent manner. (**A-B**) Kidney pictures and Periodic-acid Schiff’(PAS) staining (A) and quantification (B) of kidney weight to body weight ratio (KW/BW) of obstructed and non-obstructed kidneys control, *Pkd1*^Δtub^ and *Pkd1*^Δtub^; *Kif3a*^Δtub^ mice four days after unilateral ureteral obstruction (UUO) performed at 8 weeks of age. (**C-D**) Kidney pictures and PAS staining (C) and quantification (D) of KW/BW of kidneys from *Kif3a*^Δtub^ and *Pkd1*^Δtub^; *Kif3a*^Δtub^ fourteen days after UUO performed at 8 weeks of age (D). Single *Kif3a* mutant mice (*Kif3a*^Δtub^ mice) are presented instead of control to show epistasis of *Kif3a* over *Pkd1* inactivation. Scale bars: 1 mm (kidney pictures) and 0.1 mm (PAS). Each pair of linked symbols (dot and square) represents the obstructed (square) and non-obstructed (dot) kidneys of an individual female mouse. Scale bars: 10 µm. Two-way ANOVA, P-value for obstruction (grey; *P_obs_*), genotype (brown; *P_gen_*) and their interaction (red; *P_int_*), followed by Tukey-Kramer test: *P<0.05, **P<0.01, ***P<0.001, ****P<0.0001 or the indicated P-value.

**Figure 8.**
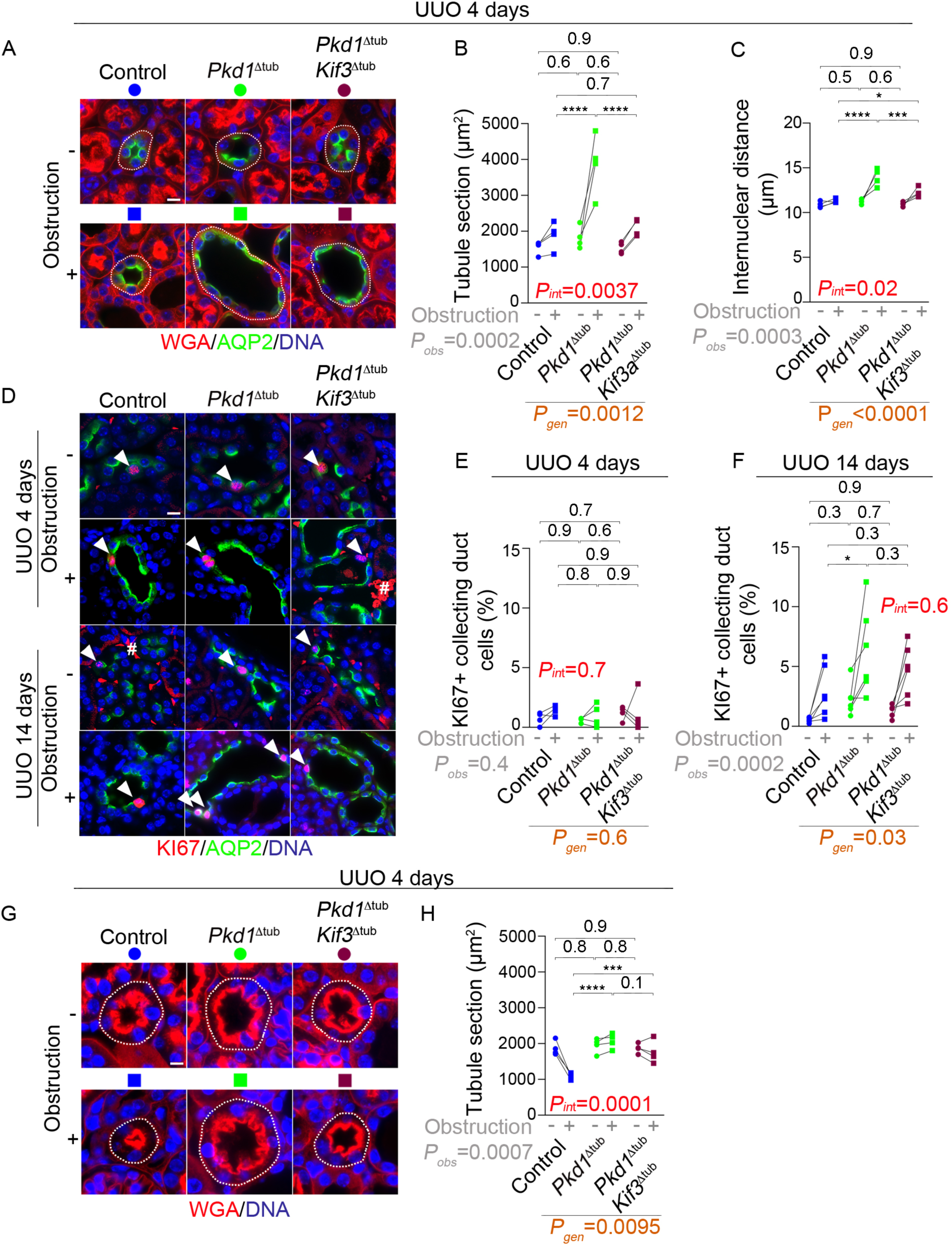
Accelerated cystogenesis after obstruction correlates with tubule distension and not proliferation. (**A-C**) fluorescent labelling of DNA, wheat germ agglutinin (WGA, which stains the brush border of proximal tubule [PT] and all basement membranes) and aquaporin 2 (AQP2, a collecting duct [CD] marker) and quantification of CD mean tubule cross-sectional area (B) and mean internuclear distance (C) in kidneys from control, *Pkd1*^Δtub^ and *Pkd1*^Δtub^; *Kif3a*^Δtub^ four days after unilateral ureteral obstruction (UUO) performed at 8 weeks of age. (**D-F**) fluorescent labelling of DNA, KI67 and AQP2 (D) and quantification (E-F) of KI67 staining (white arrowhead; # indicates non-specific red blood cells autofluorescence) in CD of kidneys from control, *Pkd1*^Δtub^ and *Pkd1*^Δtub^; *Kif3a*^Δtub^ four (E) and fourteen (F) days after UUO. (**G-H**) Representative images (G) and quantification of the PT cross-sectional area (H) in kidneys from control, *Pkd1*^Δtub^ and *Pkd1*^Δtub^; *Kif3a*^Δtub^ four days after UUO. Dashed lines in images underline tubule perimeters. Scale bars: 10 µm. Each pair of linked symbols (dot and square) represents the obstructed (square) and non-obstructed (dot) kidneys of an individual female mouse. Two-way ANOVA, P value for obstruction (grey; *P_obs_*), genotype (brown; *P_gen_*) and their interaction (red; *P_int_*), followed by Tukey-Kramer test: *P<0.05, **P<0.01, ***P<0.001, ****P<0.0001 or the indicated P-value.

In summary, the cystogenic response to obstruction corresponded with the observed effect of PC1 and cilia on TBM structure and mechanics: PC1-deficient CD and DCT have a thinner TBM and higher distensibility than PT and correspondingly dilate more strongly, precipitating cystogenesis. More generally, our results support the idea that PC1 and cilia govern tubule response to obstruction.

### Transient obstruction is sufficient to precipitate irreversible cystogenesis

Previous reports showed that only a minority of PKD cysts are obstructed at any given time^21,34^, suggesting that permanent tubule obstruction is not a frequent driver of cystogenesis. To study the effect of a transient obstruction on cyst formation, we used a reversible UUO model (R-UUO): *Pkd1*^Δtub^ mice were submitted to a ligation of the distal part of the ureter, followed by a surgical reversion of obstruction or a sham operation 4 days after UUO and the kidneys were analysed at 14 days (*i.e.*, 10 days after UUO reversion; **Figure 9A**). Remarkably, surgical reversion of obstruction after 4 days failed to reduce cyst growth at 14 days (**Figure 9B-D**), demonstrating that transient obstruction is sufficient to trigger an irreversible cystogenic program.

**Figure 9.**
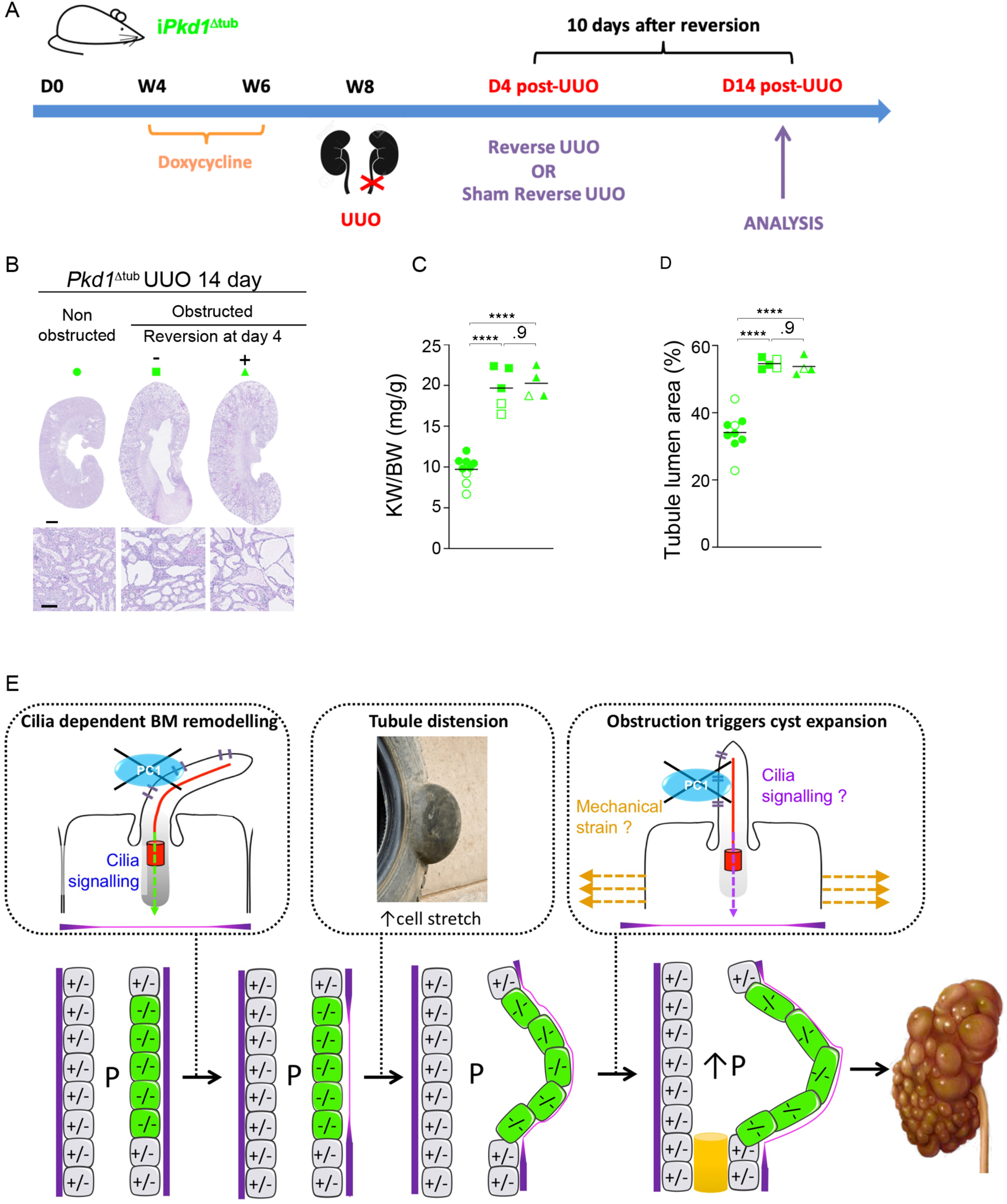
Transient obstruction triggers irreversible cystogenesis in *Pkd1* mutant mice. (**A**) Scheme of the protocol of reversible unilateral ureteral obstruction (UUO). Left kidney of 8-week-old *Pkd1* mutant mice was submitted to UUO, followed by a release of the obstruction (Reverse UUO) or a Sham operation (Sham Reverse UUO) 4 days after UUO, and mice were sacrificed 10 days after the second surgery. (**B-D**) Periodic-acid Schiff’s staining (B) and quantification of kidney weight to body weight ratio (KW/BW; C) and tubule dilations (D) of the kidneys from control or *Pkd1* mutant mice submitted to Reverse UUO or Sham Reverse-UUO. Scale bars: 2 mm (up) and 0.1 mm (down). Each symbol represents an individual female (open symbol) or male (closed symbol) mouse kidney. One-way ANOVA followed by Tukey-Kramer test: ****P<0.0001 or the indicated P-value. (**E**) Proposed “tire bulge” model for cyst formation in ADPKD: Polycystin-1 loss triggers cilia dependent remodelling of basement membrane (BM). BM remodelling in turn promotes tubule distension, which is exacerbated by transmural hydrostatic pressure (P). Transient tubule obstruction increases luminal pressure, tubule dilation and tubular cell stretch, precipitating cystogenesis.

Altogether, these results for the first time establish a TBM-centred model of cystogenesis in which *Pkd1* loss drives cilia-dependent remodelling of TBM shape and mechanics, which, in conjunction with increased luminal pressure, produce tubule dilation in a manner resembling the formation of a tire bulge (**Figure 9E**). In this condition of increased tension, obstruction precipitates cystogenesis independent of cell proliferation.

## DISCUSSION

ADPKD is the most common of the ciliopathies (*i.e.*, diseases caused by mutations in ciliary genes). While the genetics are well understood, its main feature, namely cyst formation, has not been mechanistically elucidated. We now demonstrate that cilia and PC1 regulate the composition of TBM and consequently its biomechanical properties. We show that disturbance of TBM shape and biomechanics is at the source of tubular dilation and cyst formation, defining a novel cystogenic mechanism underlying ADPKD.

Formerly, TBM remodelling of cyst epithelia has been considered a secondary “passive” mechanism^35^. Other reports have concluded that ECM stiffening results in cell proliferation *via* mechanosensitive YAP/TAZ activation, thus promoting cyst growth^29,36^, and replicating mechanisms of cell growth in cancer^37^. Instead, our data demonstrate that the TBM is altered as a direct consequence of *Pkd1* inactivation, and that it is present at the very beginning of distal tubule dilation. The remodelling of the TBM is characterized by thinning and an increase in HS, which is predicted to decrease TBM stiffness. Indeed, examining tubule-pressure curves of isolated tubules revealed a strong correlation between TBM thinning, tubule distension *in vivo* and increased tubule distensibility *ex vivo*. In line with a direct role of ECM mechanics in ADPKD, we show that increasing ECM deformability facilitates *Pkd1*^-/-^ tubule-on-chip dilation and cystogenesis in *Pkd1*^Δtub^ mice. Our data strongly suggest that TBM remodelling upon *Pkd1* inactivation and subsequent alteration in tubule mechanics play a critical role in cyst formation. This finding explains published observations from patients: various collagen mutations have been associated with the formation of renal cysts^38^ and individuals with combined *COL4A1* and *PKD2* mutations were observed to have an accelerated ADPKD course^39^.

We find that the TBM remodelling in PC1 deficiency requires cilia. Two genetic strategies preventing cilia formation, by targeting the ciliary motor subcomplex *Kif3a* and the intra-flagellar transport protein *Ift20*, consistently prevented TBM remodelling and tubule distension in PC1-deficient kidneys. These results provide the first *in vivo* evidence that cilia regulate TBM structure and composition and further link TBM remodelling with cysts formation. Altered TBM regulation may be involved in other ciliopathies. This is the case for nephronophthisis in which kidney fibrosis is associated with characteristic but unexplained TBM modifications^40^. Regulation of ECM deposition by cilia is established in mesenchymal cells such as chondrocytes in which cilia are buried into ECM and monitor its characteristics^41^. In contrast, in the kidney tubule, cilia protrude into the lumen and are believed to sense mechanical cues generated by the urine flow, possibly through PC1/PC2 complex^42,43^. Interestingly, the TBM seems to be adapted to the hydrodynamic constraints to which tubules are exposed. For instance, TBM thickness decreases along the tubule as does hydrostatic pressure^44^. Similarly, glomerular hyperfiltration, which occurs in diabetic nephropathy or after nephron reduction, is associated with TBM thickening. Thus, our findings raise the possibility that primary cilia are involved in matching TBM mechanics with the mechanical constraints imposed by the urine flow. Besides the kidney, BM remodelling plays important roles in multiple contexts including development and cancer^45^, as does the primary cilium. The connection between cilia and BM may have implications beyond the kidney field.

This study brings novel insights regarding cell proliferation in ADPKD. Abnormal cell proliferation has been shown to be an integral part of ADPKD and is considered an important driving force for cyst growth^14,46^. We find that the timing of cell proliferation as a driver of cyst growth occurs later than previously thought. During early cystogenesis our results pinpoint TBM remodelling as the cause. At first TBM remodelling is associated with early distal nephron distension independently of cell proliferation. At a later stage proliferation and tubule distension occur concurrently. UUO experiments allowed us to dissociate proliferation, distension and cystogenesis further: obstruction resulted in an increase in tubular cell proliferation across all genotypes. However, the initial tubule dilation only progressed in PC1-deficient DCT and CD, where it coincides with TBM remodelling and excessive cell stretching. Mechanistically, by increasing the area available for the intercalation of daughter cells in the epithelial layer, TBM remodelling may both reduce cell crowding, which causes cell loss through extrusion and/or apoptosis, and increase cell stretching, which accelerates G2/M transition^47^.

Our results bring further insight into the role of obstruction in ADPKD. In the human ADPKD kidney obstruction occurs mostly through cyst enlargement and possibly due to crystal formation. Crystals were shown to accelerate cystogenesis in animal models of PKD^48^. The cyst-promoting effect of obstruction may explain the ‘snowball effect’ of cystogenesis, an observation that small cysts preferentially occur in the neighbourhood of large cysts^49^. Our results provide strong evidence that obstruction, even if transient, is a potent driver of cyst formation. Tubular cell injury has been proposed as a common mechanism responsible for accelerated cyst formation in the setting of ischemia-reperfusion or nephrotoxic drugs^48–50^. Yet, by showing that obstruction promotes cyst formation selectively in the distal nephron, our results pinpoint excessive mechanical strain, which is restricted to this segment, rather than injury, which also affect PT, as the main factor precipitating cyst formation.

Importantly, our animal models recapitulate a finding from human ADPKD kidneys, *i.e.* that cysts occur mostly in distal nephron segments, and provide a mechanical explanation to it: CD and DCT (i) have thinner and HS enriched TBM, (ii) precociously activate genes involved in TBM remodelling in response to PC1 loss and (iii) undergo early distension with altered biomechanics, which culminates in excessive cell stretching and accelerated cytogenesis after obstruction.

In summary, we show that PC1 and cilia regulate the composition and mechanical properties of the TBM. Our findings reveal a novel function of primary cilia and for the first time provide a biomechanical model to explain cyst formation in ADPKD.

## METHODS

### Mice

Mice were housed at constant ambient temperature in a free-pathogen facility on a 12-hour day/night cycle and were fed *ad libitum*. Breeding and genotyping were performed according to standard procedures. All mice were on a C57BL/6 (mixed J/N) background. *Pkd1*^flox/flox^ mice (B6.129S4-Pkd1^tm2Ggg/J^, Jackson Laboratory), *Kif3a*^flox/flox^ mice (Kif3a^tm1Gsn^, C57BL/6 genetic background, kindly provided by Peter Igarashi, Stony Brook University), *Ift20*^flox/flox^ mice (Ift20^tm1.1Gjp^, kindly provided by Gregory Pazour, UMass Chan Medical School) were crossed to Pax8rtTA (Tg[Pax8-rtTA2S*M2]1Koes, Jackson Laboratory) and TetOCre (Tg[tetO-cre]1Jaw, Jackson laboratory) mice to create mice with inducible tubule-specific *Pkd1* knockout (named as *Pkd1^Δtub^*), double *Pkd1*; *Kif3a* knockout (named as *Pkd1^Δtub^*; *Kif3a^Δtub^*) and double *Pkd1*; *Ift20* knockout (named as *Pkd1^Δtub^*; *Ift20^Δtub^*). *Pxdn*-deficient mice (referred to as *Pxdn*^-/-^, C57BL/6 genetic background) were kindly provided by Miklos Geiszt (Semmelweis University)^51^ and were crossed with *Pkd1^Δtub^* mice to generate *Pkd1^Δtub^* mice with or without constitutive *Pxdn* inactivation (named as *Pkd1^Δtub^*; *Pxdn*^-/-^).

All animals received doxycycline hyclate (Abcam, ab141091, 2 mg/mL) in drinking water supplemented with 5% sucrose and protected from light, from postnatal day 28 to 42 to induce the inactivation of floxed alleles. Animals lacking either TetOCre or Pax8rtTA were used as controls. Both gender were used in experiments.

### Animal procedures

Euthanasia of mice consisted in an intraperitoneal injection of anesthetizing solution (ketamin 160 mg kg^-1^ and xylazin 8 mg kg^-1^) followed with cervical dislocation once the depth of anaesthesia was confirmed.

Unilateral ureteral obstruction (UUO) was performed in 8-week-old animals as previously described^52^. Briefly, mice received an intraperitoneal injection of anesthetizing solution (ketamin 160 mg.kg^-1^ and xylazin 8 mg.kg^-1^) associated with intraperitoneal injection of buprenorphine (0.1 mg.kg^-1^) to prevent postoperative pain. Under optical microscope, a left-side laparotomy was performed. The left ureter was exposed and ligated with a nonabsorbable 6/0 black braided silk suture. Reversion of ureteral obstruction was performed as previously described^53^. A midline laparotomy was made to expose the left ureter. Firstly, the left ureter was ligated twice with a nonabsorbable 6/0 black braided silk suture. A longitudinally split sterile silicone tube (soft walled silicone plastic tubing) was then gently placed around the ureter. At last, a ligature around the tube was made twice to anchor and maintain the tube closed before closing the incision. Four days after the first surgery of UUO, the abdominal wall was re-opened. The silicon tube was removed and the left ureter was cut between the two sutures. The superior part of the ureter was then re-implanted within the bladder. Ureter section and reimplantation steps were skipped for sham reimplantation surgery.

### Standard morphological analysis

For histopathological analysis, mouse kidneys were fixed in 4% paraformaldehyde for 24 hours at 4°C and embedded in paraffin. Four micrometres thick sections were stained with Periodic acid-Schiff (PAS). PAS-stained, full-size kidney sections were imaged using a whole-slide scanner Nanozoomer 2.0 (Hamamatsu) equipped with a 20X /0.75 NA objective coupled to NDPview software (Hamamatsu). Quantification of tubule dilation was performed on PAS-stained whole kidney sections with ImageJ2 software Version 2.9.0/1.53t, using NDPITools Plugin (version 1.8.3)^54^. Briefly, this plugin automatically split the original PAS kidney sections into several images at magnification 20X. Cortical fields were then manually selected across the renal cortex on each image. The tubular lumen surface was automatically measured in each extracted cortical field, as previously described^55^, and the results were finally expressed as percentages of total cortical tubular lumen area of the selected fields.

### Fluorescent kidney staining

Four micrometres thick sections of paraffin-embedded kidneys were submitted to deparaffinization and hydration. Kidney sections were stained with the molecules listed in **Supplementary table 1.** We used distinct antigen retrieval: incubation in 10mM Tris Base, 1mM EDTA Solution, 0.05% Tween 20, pH 9.0 or citrate buffer pH6 (Zytomed, ZUC028) for 15 minutes in high pressure condition in a TintoRetriever® pressure cooker (BioSB) for *Wheat Germ Agglutinin* (WGA), Aquaporin 2 (AQP2), Calbindin (CALB), KI67 and Proliferating Cell Nuclear Antigen (PCNA) labelling or in 50 mM Tris base, 1 mM EDTA solution, 0.01% proteinase K (Roche, 3115836001) for 2 minutes at room temperature, followed by avidin/biotin blocking (Vector Laboratories, SP-2001) for collagen IV, laminin and HSPG labelling. Then, kidney sections were incubated with primary antibody overnight at 4°C, followed with the appropriate Alexa Fluor-conjugated secondary antibody.

Images were acquired using the Zeiss Apotome 2 CO2 at the original magnification of 40X or Zeiss Spinning Disk L3 at the original magnification of 63X, and processed with ImageJ2 software.

### BrdU incorporation assay

Six hours following UUO, control, *Pkd1^Δtub^* and *Pkd1^Δtub^*; *Kif3a^Δtub^* mice received a single intraperitoneal injection of BrdU (100 mg/kg) and were sacrificed 18 hours after BrdU injection (*i.e.*, 24 hours after UUO). Kidney sections were immersed in Tris-EDTA based solution, for 15 minutes at 95°C and stained with the corresponding anti-BrdU antibody (**Supplementary table 1**).

### Quantification of tubular cell proliferation

The tubular cell proliferation index was calculated as the number of KI67, PCNA or BrDU positive nuclei over the total number of tubular nuclei for each nephron segment, using the “Cell Counter” plugin of ImageJ software. It was scored in at least 10 randomly selected fields per kidney section.

### Quantitative tubule morphometric analysis

The tubule cross-sectional area (expressed in μm^2^) was automatically quantified with ImageJ2 software using U-Net Segmentation Plugin^56^. Briefly, the outer tubular surface of proximal tubules (PT; brush border stained with WGA), distal convoluted tubules (DCT; stained with CALB) and collecting ducts (CD; stained with AQP2) were annotated on a few pictures of labelled kidneys. We use an Ubuntu server 18.04 with FIJI, a distribution of ImageJ, and the U-Net Segmentation plugin (https://sites.imagej.net/Falk/) that interfaces artificial neural networks. A pre-trained model (2d_cell_net_v0) was used and adjusted with the plugin option ‘U-Net Finetuning’ : manual annotation on personal pictures: extpct 129, extdistal 76, glom 5. The model was applied to segment tubules in large regions of the cortex.

Quantification of internuclear distance was performed manually on each image using the *Line Selection Tool* in ImageJ2 software, as the distance (expressed in μm) between the centre of two adjacent nuclei along the tubular wall. It was measure for each specifically stained nephron segment of interest: the proximal tubule, the distal convoluted tubule and the collecting duct. To facilitate the comparison of the morphometric data obtained in 8 and 12 weeks-old animals, the cross-sectional area and internuclear distance measured at 12 weeks were divided by the mean observed in control mice at 12 weeks and multiplied by the mean value observed in control mice at 8 weeks.

### Measurement of TBM components staining intensity

Fluorescence intensity of TBM components (HSPG, collagen IV, laminin) was automatically quantified using a specific ImageJ2 macro, named *“Do_ROI_dilate_manager”*. Briefly, it consisted in surrounding the basement membranes stained with WGA of each tubule of interest on immunofluorescence kidney sections, using the *Area Selection Tool* in Image J2. Each selected area was extracted as a ROI (Region Of Interest), then listed in the ROI Manager and individually dilated by the macro to capture all TBM signals. For each tubule, fluorescence intensity of TBM components was assessed by the *Mean Gray Value*. The mean fluorescence intensity per mouse was then calculated as the mean of the mean gray values of individual tubules.

### Transmission electron microscopy

Five-mm cortical slices of each kidney were fixed with 2.5% glutaraldehyde in 0.1 M sodium cacodylate buffer prepared at pH 7.4 (Euromedex, 12300-25) for 24 hours at 4°C. After rinsing with 0.2 M sodium cacodylate buffer, the kidneys samples were maintained in the same buffer at 4°C. Subsequently, they were post-fixed in 1% OsO4, dehydrated, and embedded in epoxy resin blocks. Blocks were then cut with an UC7 ultramicrotome (Leica, Leica Microsystemes SAS). Finally, 70 nm-thick sections were recovered on copper and visualized with Hitachi HT7700 electron microscope. Pictures (2048x2048 pixels) of PTs, DCTs and CDs were imaged with an AMT41B and processed using ImageJ2 (Version 2.9.0/1.53t). TEM acquisition was performed by the Imaging Facility of Hôpital Tenon (INSERM UMRS_S1155). The mean basement membrane thickness was measured in a blind fashion using at least 3 measurements per tubule.

### Measurement of pression-diameter curves in isolated perfused tubule

Kidneys from control and *Pkd1*^Δtub^ mice were collected at 8 and 12 weeks and were cut into slices. PT and CD were then dissected without collagenase digestion, to preserve TBM mechanics. Each isolated tubule was transferred to a 37°C controlled-bath chamber under an inverted microscope (Axiovert 100, Carl Zeiss) and mounted between two concentric glass pipettes, as previously described^57^. The perfusion and bath solutions were composed of 144 mM NaCl, 5.5 mM glucose, 5 mM alanine, 10 mM HEPES, 1.5 mM CaCl_2_, 1.2 mM MgSO_4_, 2 mM K_2_HPO_4_, pH 7.4. The TBM of isolated tubule was stained for 5 minutes by adding fluorescein-labeled WGA within the bath chamber (**Supplementary table 1**). A solution column connected to the perfusion pipette was vertically displaced, allowing for progressive increment of transmural pressure delivered to the tubule lumen, from 0 to 42 cmH_2_0. At each pressure point, the outer tubular diameter was measured (mean of 10 measurements per tubule).

### Microdissection of tubule segments

Deeply anesthetized mice were placed under binocular microscope and submitted to a laparotomy, followed by an incision of subcutaneous and muscular layers to gain access to the peritoneal cavity. A first vascular clamp was applied to the abdominal aorta to prevent bleeding during aortic incision. Three other ligatures were placed on the abdominal aorta at different locations before aortic incision: at the distal end to facilitate microperfusion, at the mid part to prevent catheter displacement and at the proximal end before the bifurcation of the left renal artery to maximize the perfusion of the left kidney. An incision was then performed in the wall of the abdominal aorta and a catheter linked to a syringe was inserted in the lumen towards the left renal artery. The left kidney was rinsed 5 ml of dissection buffer [Hank’s medium without phenol (L0612-500, Eurobio) complemented with 1 M MgCl_2_, 3 M sodium acetate, 100 mM sodium pyruvate, 200 mM glutamine, 1 M HEPES, 0.1% BSA, pH 7.4] and perfused with the same medium containing 120 μg/ml of Liberase (Roche, 05401127001). Thin kidney slices were cut along the corticomedullary axis and incubated at 30°C for 25 minutes in dissection buffer containing 40 μg/ml of Liberase. Kidney slices were then transferred in Petri dishes containing dissection buffer. The PTs and the CDs were microdissected under binocular magnifier according to morphological and topographic criteria, using fine needles. Approximately 150 PTs and 150 CDs were isolated from each kidney.

### RNA extraction and RNA-seq pre-processing and analysis

Total RNAs were extracted and purified from the microdissected PTs and CDs using RNeasy Micro Kit (Qiagen; 74004) according to the manufacturer’s protocol. RNA quality was verified by capillary electrophoreses using Fragment Analyzer^TM^.

Bulk mRNA-Seq libraries were prepared by the Imagine Genomic Core Facility (Paris, France) starting from 7ng of total RNA, using the NEBNext® Single Cell/Low Input RNA Library Prep Kit according to manufacturer’s guidelines. This kit generates mRNA-Seq libraries from the PolyA+fraction of the total RNA by template switch reverse transcription. The cDNA are amplified twice and the final mRNAseq libraries are not ‘oriented’ or ‘stranded’ (meaning that the information about the transcribed DNA strand is not preserved during the library preparation). The equimolar pool of libraries, assessed by Q-PCR KAPA Library Quantification kit (Roche) and with a run test using the iSeq100 (Illumina) was sequenced on a NovaSeq6000 (Illumina, S2 FlowCells, paired-end 100+100 bases, ∼50 millions of reads/clusters produced per library). After the demultiplexing and before the mapping, the reads were trimmed to remove the adaptor sequences from the first amplification. Reads were mapped to the GRCm38 primary mouse genome assembly with STAR^58^. Counts table was generated with feature Counts from subread-1.6.4 package^59^. Normalization and differential gene expression DGE analyses were performed with R package DESeq2 v1.32.0 (doi:10.1186/s13059-014-0550-8), and Benjamini-Hochberg correction was applied with threshold for significance set at adjusted P-values < 0.05.

Based on data from bulk RNASeq generated in tubules from either *Pkd1* mutant or control mice, we identified the genes that are differentially expressed in collecting ducts and proximal tubules by using the package Deseq2 . Gene set enrichment analyses were performed on GSEA v4.3.2 software (Broad Institute).

### Analysis of BM remodelling signature expression in human single nuclei RNAseq dataset

We downloaded a single nuclei dataset from human PKD^27^ on the Gene Expression Omnibus platform (GSE185948). After using a mouse-human ortholog table, we calculated a z-score of the signature for each cell from the dataset. Violin plot according to the cell type was then generated using the VlnPlot function from Seurat. We used the cell type annotations identified in the original article^27^. UMAP expression plots for selected genes were generated thanks to KIT interactive web site (http://humphreyslab.com/SingleCell/)^60^.

### Tubule-on-chip fabrication and culture conditions

As previously described^61^, the chip consists of a PDMS scaffold of 3 PDMS layers bonded to a glass slide. One part of the scaffold features the microfluidic channels. For this part, a PDMS (Curing Agent to PDMS weight ratio of 1:10) was poured on a mold obtained with photolytography. This part lays on a flat PDMS membrane which elevates the channels from the glass slide. This flat PDMS layer was obtained by spincoating PDMS 1:10. On top, another flat 500-µm thick PDMS membrane closes the chip. The PDMS scaffold surface was activated oxygen plasma and then treated with a 5% (3-aminopropyl) triethoxysilane (Sigma-Aldrich) solution in methanol for 45 min, rinsed thoroughly in methanol in an ultrasonic bath then in a 2.5% glutaraldehyde (Sigma-Aldrich) solution in ddH20 for 45min and rinsed thoroughly in ddH20 in an ultrasonic bath then at room temperature overnight. Then, 75 µm wide tungsten wires were inserted into the scaffold, which was sterilized with UV-Ozone. A 6g/L or 9.5g/L collagen I gel was prepared on ice by neutralizing high concentration collagen gel from rat tail with NaOH and diluted in PBS with 10% of total volume with PBS 10X. The gel was injected in the collagen chambers, still on ice. The collagen was set at 37°C for 2 hours. Chips were then stored in PBS at 4°C up to 2 weeks before use.

The day before seeding, wires were partially removed to form the channels and a laminin 50µg/mL solution was injected in the chip using a pressure controller and PTFE tubing followed by 1 hour-incubation at 37°C. For the seeding, we used an edited clonal mIMCD-3 cell line (immortalized cells from murine inner medullary collecting duct), deleted for *Pkd1* gene using CRISPR-based genome editing, kindly provided by Michael Köttgen^31^. These cells were detached from flask upon confluency with trypsin, counted and resuspended at 10 million cells/mL in DMEM/F-12 medium with 10% FBS. The cell suspension was then injected into collagen channels with a pressure controller via PTFE tubing until the channels are filled with cells. Then tungsten wires were inserted back into the channels to form a hollow tube. Chips were incubated in DMEM/F-12 medium with 10% FBS for 2 to 3 hours to let cells adhere. Then tungsten wires were removed to obtain a confluent mIMCD-3 layer. Chips were then cultured in 6-well plates in humid atmosphere at 37°C for up to 2 weeks in DMEM/F-12 medium with 10% FBS. Medium was changed every two days. Chips were then washed with PBS twice before being fixed in a 4% paraformaldehyde solution in PBS for 45 min. Chips were then rinsed twice in PBS and stored in PBS at 4°C before immunostaining.

### Tubule-on-chip immunolabelling

The cells were permeabilized by submerging the chip into PBS + 0.3% TritonX100 for 30 minutes, and rinsed three times for 30 minutes with PBS-BSA 2%. The chip was submerged in PBS-BSA 2% with 0.1% Tween20 and 2% Goat Serum for saturation. A 100-150 µL drop of 1/200 antibody, 2% BSA, 1% Tween20 in PBS was delicately placed on the chip so that it does not fall. Incubation was carried out for 24 hours in a humid environment at room temperature. The chip was rinsed 3 times for 30 minutes in PBS-BSA 2%. A 100 µL drop of a solution containing 1 µg/ml Hoechst, 0.25 µg/ml fluorescent phalloidin, rabbit anti-KI67, (Abcam, ab16667, 1:100), 2% BSA, 1% Tween-20 in PBS was delicately placed to prevent spilling. Incubation was carried out for 24 hours in a humid environment at room temperature.

### Data Analysis and Statistics

Data were expressed as means. Differences between the experimental groups were evaluated using one or two-way ANOVA followed, when significant (P < 0.05), by the Tukey-Kramer test. When only two groups were compared, unpaired t-test or Mann-Whitney test was used as appropriate. No randomization was performed. Quantification of TBM thickness and morphometric analysis were performed in a blind fashion. Linear regression was also used to correlate tubule section and internuclear distance in the CD and the DCT at 8 weeks. The statistical analysis was performed using GraphPad Prism V10 Software.

### Data Sharing statement

Upon publication the transcriptomic datasets that we generated will be deposited in Gene Expression Omnibus platform and made fully available.

All the mice models used in this study can be obtained upon request to the corresponding author, with the agreement of the scientist who generated the initial transgenic line.

### Ethics

All mice experiments were conducted in accordance to the guidelines of the French government animal welfare policy. Animal procedures were approved by the ethical committee of the Ministère de l’Enseignement Supérieur, de la Recherche et de l’Innovation (APAFIS agreements #201907041733347, #2020090715389782, #2020111915598007).

## ACKNOWLEDGMENTS

We thank the animal facility (LEAT, SFR Necker, INSERM US24, Paris, France), the histology facility (SFR Necker, INSERM US24, Paris, France), the imaging platform (SFR Necker, INSERM US24, Paris, France) and the genomic core facility (SFR Necker, INSERM US24, Paris, France) for technical assistance. This work benefited from the technical contribution of the joint service unit CNRS UAR 3750). We additionally thank Gregory Pazour, Peter Igarashi and Miklos Geist for providing transgenic mice and Benjamin Humphrey for allowing us to use his Kidney Interactive Transcriptomics interface and human snRNAseq data. We further thank Prof. Vincent Durlach for his support.

## FUNDING

This work has received support from the *Agence Nationale de la Recherche* (ANR-19-CE14-0016 Homeocyst, recipient F. Bienaimé). This work was supported by doctoral grants from AMX (“*Ministère de la Recherche et de l’Enseignement supérieur”*, France ; recipient B. Lapin), the ‘*Fondation pour la Recherche Médicale*’ (FRM ; recipients M. Mazloum: FDM201906008513 provided by the Durlach foundation, and B. Lapin: program FDT) and King Abdulaziz University (Jeddah, Saudi Arabia ; recipient R. Alghamdi), a postdoctoral grant from the FRM (recipient A. Viau FRM ARF20150934110), an Interface contract from the *Institut National de la Santé et de la Recherche Médicale* (INSERM ; *Contrat d’interface INSERM ;* recipient F. Bienaimé). This work was also supported by the European Commission grant FET Open program (FETOPEN-01-2016-2017). This work has received the support of “*Association Polykystose France*”, provided by the “*Société Francophone de Néphrologie*”*, Dialyse et Transplantation*’ (SFNDT). This work has received the support of “*Institut Pierre-Gilles de Gennes laboratoire d’excellence*, *Investissements d’avenir*” program ANR-10-IDEX-0001-02 PSL and ANR-10-LABX-31. This work has received support from the grants ANR-11-LABX-0038, ANR-10-IDEX-0001-02. This work was supported by the “*Centre National de la Recherche Scientifique*” (CNRS), *Sorbonne Université* and *Institut Curie*. EWK is supported by DFG KU1504/8-1 and DFG KU1504/9-1.

## AUTHOR CONTRIBUTIONS

M.M, A.V, F.T and F.B designed the study; M.M, B.La, A.V, M.B, R.A, P.H, A.A, L.C, G.C, M-C.V, B.Le, S.C and F.B performed experiments; SG generated image analysis pipeline, M.M, F.B, R.A, B.La, S.D. and S.C analyzed data; M.M and F.B drafted and revised the manuscript with the help of A.V, S.C, E.W.K and F.T; all authors approved the final version of the manuscript.

## CONFLICT OF INTEREST

The authors declare that they have no conflict of interest.

## Notes

### Competing Interest Statement

The authors have declared no competing interest.

