## Supplementary table and figures for "Cilia to basement membrane signalling is a biomechanical driver of autosomal dominant polycystic kidney disease"

SUPPLEMENTARY MATERIAL TABLE OF CONTENTS

**Supplementary table 1.** List of the antibodies used in this study

**Extended data figure 1.** *Pkd1* deletion drives cilia-dependent distal tubule distension independently of cell proliferation

**Extended data figure 2.** *Pkd1* deletion induces cilia-dependent TBM remodelling

**Extended data figure 3.** *Pkd1* deletion induces cilia-dependent TBM thinning

**Extended data figure 4.** RNAseq profiling of micro-dissected proximal tubules and collecting ducts

**Extended data figure 5.** Control and *Pkd1*-deficient tubules display an elastic behavior

**Extended data figure 6**. Ureteral obstruction accelerates Polycystin-1 deficient distal tubule dilation in a cilia-dependent manner

**Extended data figure 7**. Ureteral obstruction triggers explosive cystogenesis in *Pkd1*$^{\Delta tub}$ distal tubules in a cilia-dependent manner

**Extended data figure 8.** Variation in cell proliferation does not explain cilia-dependent *Pkd1*$^{\Delta tub}$ tubule dilation after obstruction

Supplementary table 1. List of the antibodies and dyes used in this study

| Reference | Primary antibody/Dye/Lectin | Dilution | Source |
|---|---|---|---|
| sc-515770 | Mouse anti-AQP2 | 1:200 | Santa Cruz Biotechnology |
| sc-9882 | Goat anti-AQP2 | 1:200 | Santa Cruz Biotechnology |
| A7310 | Rabbit anti-AQP2 | 1:200 | Sigma-Aldrich |
| ab6326 | Rat anti-BrdU | 1:100 | Abcam |
| sc-365360 | Mouse anti-CALBINDIN | 1:200 | Santa Cruz Biotechnology |
| NB110-59981 | Rabbit anti-COL4A1 | 1:200 | Novus Biologicals |
| H3570 | Hoechst 33341 | 1:1000 | Thermo Fisher |
| A kind gift of Brigitte Lelong | Rabbit anti-HSPG | 1:100 | Makino H *et al* (1986)[71] |
| ab16667 | Rabbit anti-KI67 | 1:100 | Abcam |
| M0879 | Mouse anti-PCNA | 1:1000 | Dako |
| | Phallodin | | |
| FL-1021 | Fluorescein-labeled WGA | 1:200 | Vector Laboratories |
| RL-1022 | Rhodamine-labeled WGA | 1:200 | Vector Laboratories |

| Reference | Secondary antibody | Dilution | Source |
|---|---|---|---|
| A-21202 | Donkey anti-mouse, Alexa Fluor 488 | 1:500 | Thermo Fisher Scientific |

| | | | |
|---|---|---|---|
| A-21447 | Donkey anti-goat, Alexa Fluor 647 | 1:500 | Thermo Fisher Scientific |
| A-21206 | Donkey anti-rabbit, Alexa Fluor 555 | 1:500 | Thermo Fisher Scientific |
| A-21208 | Donkey anti-Rat, Alexa Fluor 488 | 1:500 | Thermo Fisher Scientific |

| Reference | Biotinylated antibody | Dilution | Source |
|---|---|---|---|
| RPN1004V | Biotinylated Donkey anti-rabbit | 1:200 | GE Healthcare |

| Reference | Streptavidin conjugate | Dilution | Source |
|---|---|---|---|
| S-32354 | Streptavidin conjugate, Alexa Fluor 488 | 1:500 | Thermo Fisher Scientific |
| S-32355 | Streptavidin conjugate, Alexa Fluor 555 | 1:500 | Thermo Fisher Scientific |

SUPPLEMENTARY FIGURES (WITH LEGENDS)

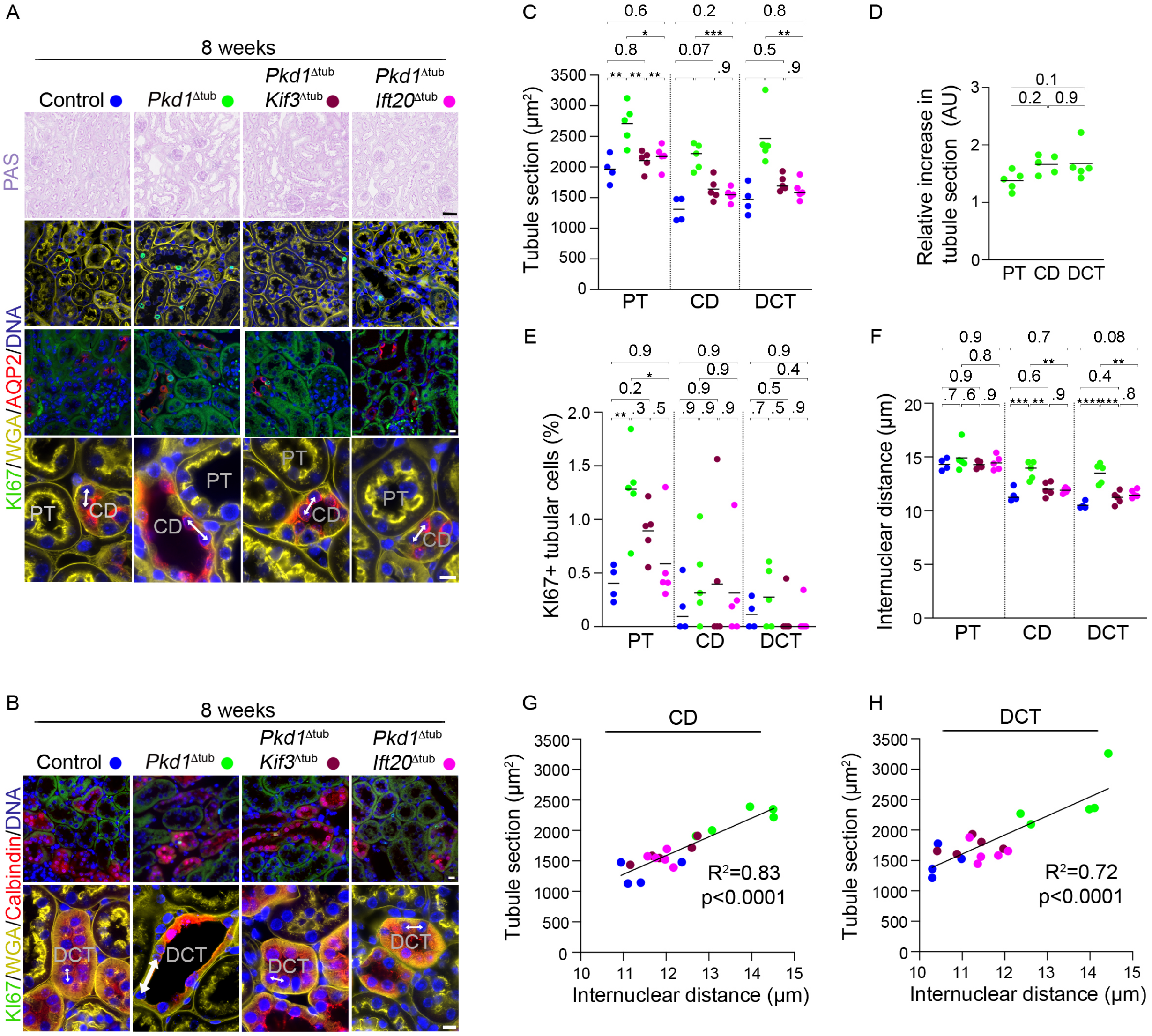

**Extended data figure 1. *Pkd1* deletion drives cilia-dependent distal tubule distension independently of cell proliferation**. (**A-B**) Periodic acid Schiff's (PAS) staining (upper panel A) and labelling of KI67 (which stains proliferating cell), DNA, wheat germ agglutinin (WGA, which stains the brush border of proximal tubule [PT] and all basement membranes), aquaporin 2 (AQP2, a collecting duct [CD] marker; lower panel A) or calbindin (a distal convoluted tubule [DCT] marker; B) of kidneys from 8-week-old control, *Pkd1*$^{\Delta tub}$, *Pkd1*$^{\Delta tub}$; *Kif3a*$^{\Delta tub}$ mice and *Pkd1*$^{\Delta tub}$; *Ift20*$^{\Delta tub}$ mice. Arrows: examples of internuclear distance measurements. Scale bars: 10 µm. (**C-F**) Quantification of mean PT, DCT and CD cross-sectional area (B), relative cross-sectional area increase in PT, DCT and CD from *Pkd1*$^{\Delta tub}$ mice (*i.e.,* normalized to the mean of PT or CD cross-sectional area of control mice; C), proliferation index (percentage of KI67+ cells; D) and internuclear distance (E) in kidneys from 8-week-old control, *Pkd1*$^{\Delta tub}$, *Pkd1*$^{\Delta tub}$; *Kif3a*$^{\Delta tub}$ and *Pkd1*$^{\Delta tub}$; *Ift20*$^{\Delta tub}$ mice. (**G-H**) Linear regression of tubule cross-sectional area and internuclear distance for CD (G) and DCT (H) in the same groups of mice. Each dot represents one individual male mouse. One-way ANOVA followed by Tukey-Kramer test: *P<0.05, **P<0.01, ***P<0.001, ****P<0.01 or the indicated P-value.

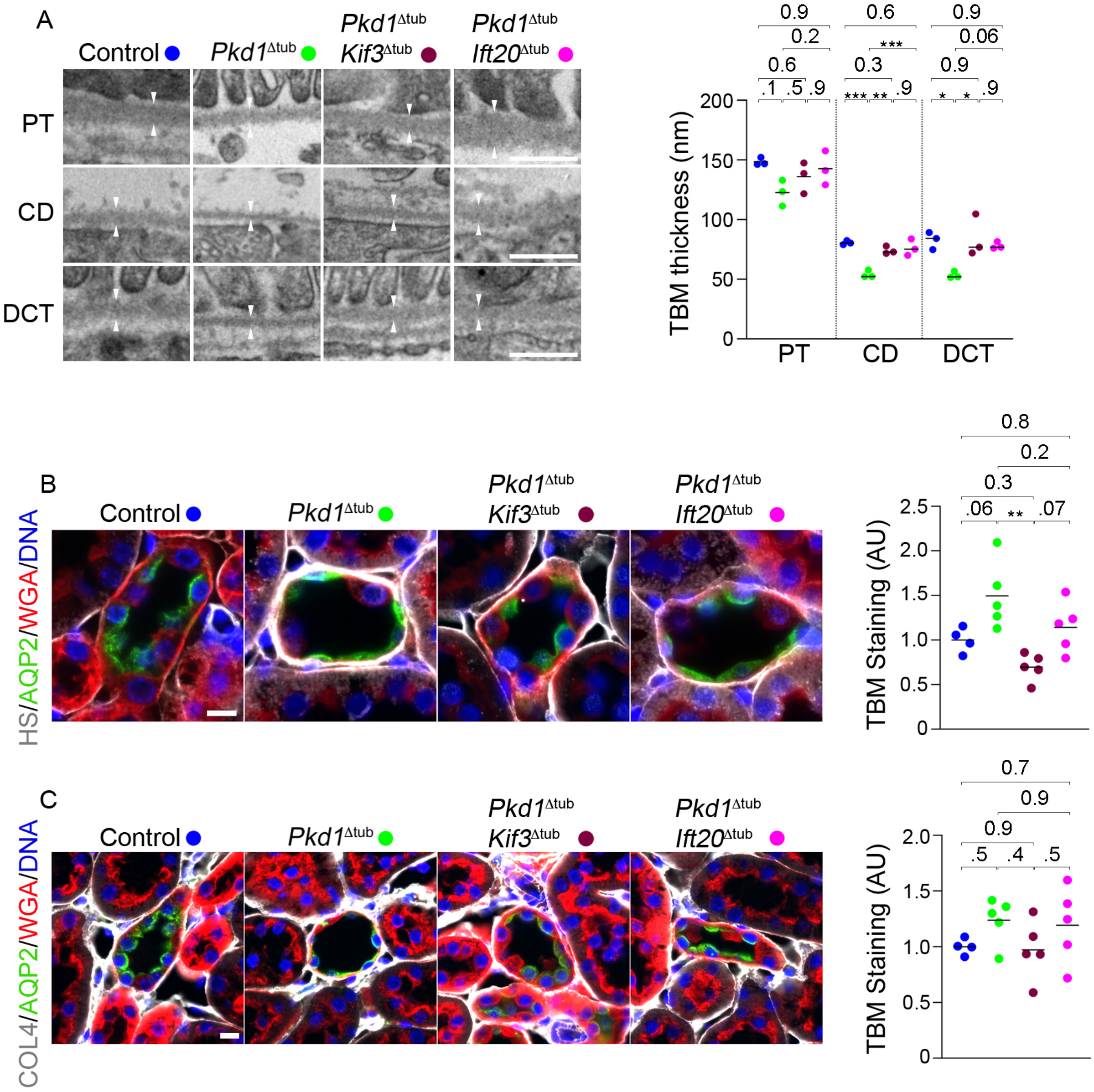


**Extended data figure 2. *Pkd1* deletion induces cilia-dependent TBM remodelling. (A)** Transmission electron microscopy and quantification of basement membrane (TBM) thickness in proximal tubule (PT), collecting duct (CD) and distal convoluted tubule (DCT) from 8-week-old control, *Pkd1*$^{\Delta tub}$, *Pkd1*$^{\Delta tub}$; *Kif3a*$^{\Delta tub}$ and *Pkd1*$^{\Delta tub}$; *Ift20*$^{\Delta tub}$ mice. Each dot represents one individual male mouse. Scale bar: 0.5 µm. (**B-C**) Staining and quantification of heparan sulfate (HS; B) and collagen IV (COL4; C) in CD of the same animals. Each dot represents one individual male mouse. Scale bars: 10 µm. One-way ANOVA followed by Tukey-Kramer test: *P<0.05, **P<0.01, *** P<0.001 or the indicated P-value. AU: arbitrary unit.

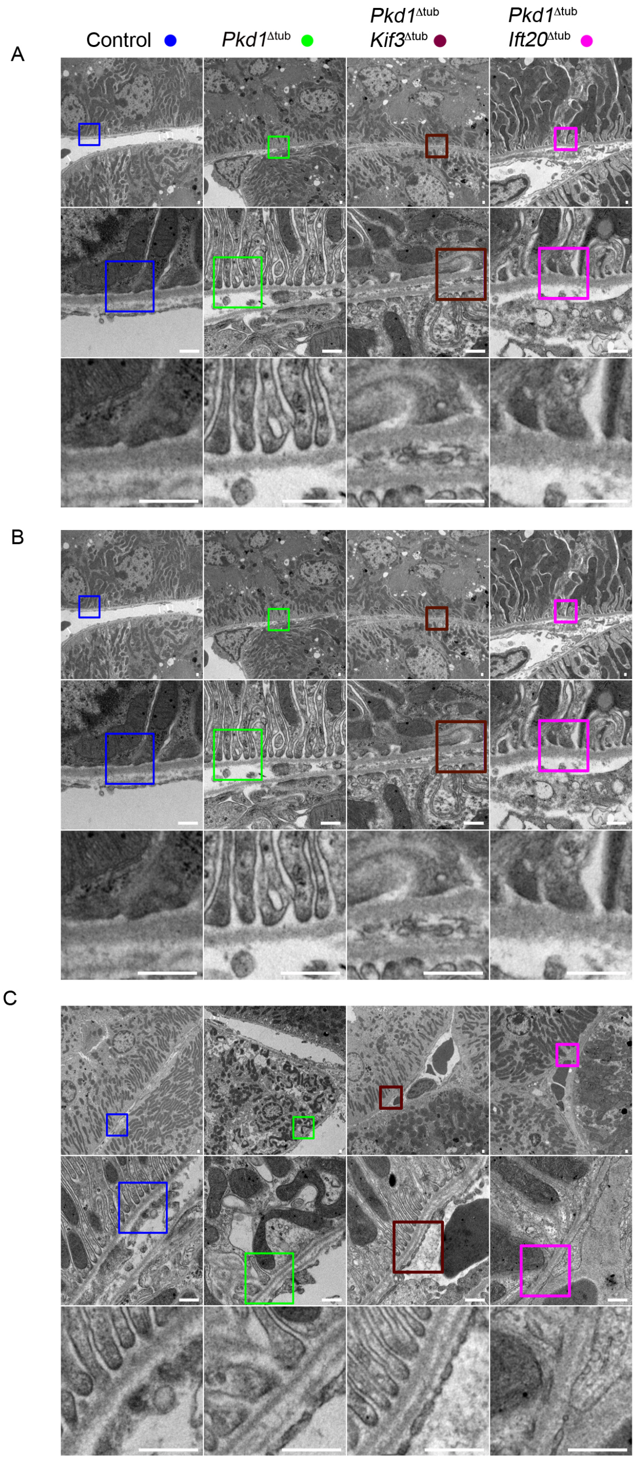


**Extended data figure 3. *Pkd1* deletion induces cilia-dependent TBM thinning.** Transmission electron microscopy of basement membrane thickness in proximal tubule (A), collecting duct (B) and distal convoluted tubule (C) from 8-week-old control, *Pkd1*$^{\Delta tub}$, *Pkd1*$^{\Delta tub}$; *Kif3a*$^{\Delta tub}$ and *Pkd1*$^{\Delta tub}$; *Ift20*$^{\Delta tub}$ mice.

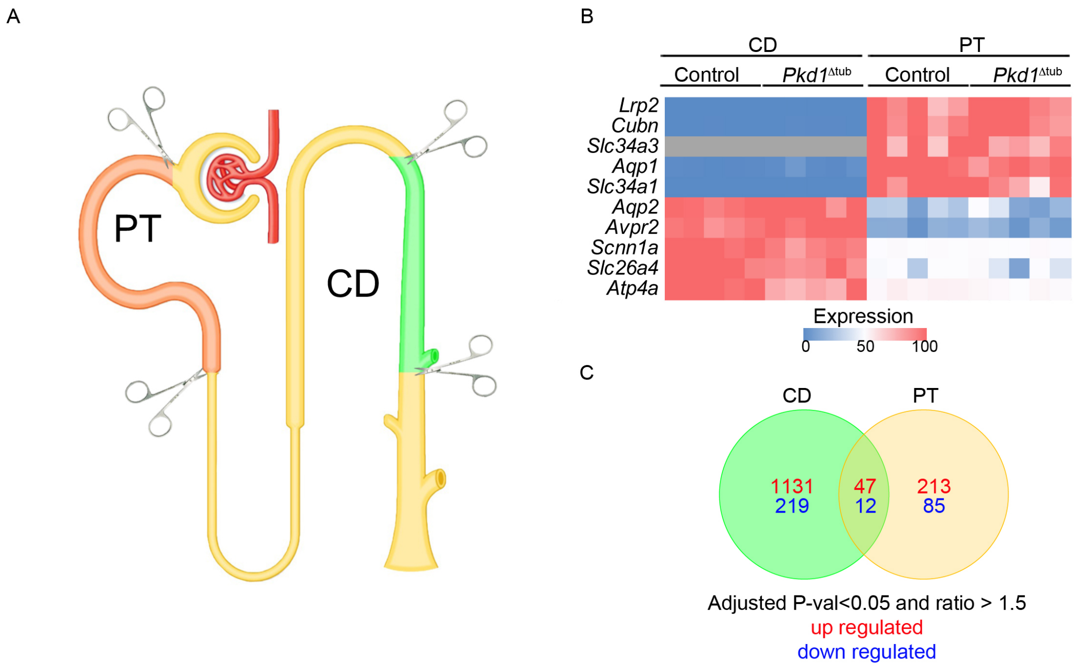


**Extended data figure 4. RNAseq profiling of micro-dissected proximal tubules and collecting ducts.** (**A**) Schematic representation of the dissected segments. (**B**) Heatmap of the relative expression of proximal tubules (PT) and collecting ducts (CD) specific genes in micro-dissected tubules. (**C**) Venn diagram showing the number of genes with significant variation between *Pkd1* mutant animals and controls in CD and PT.

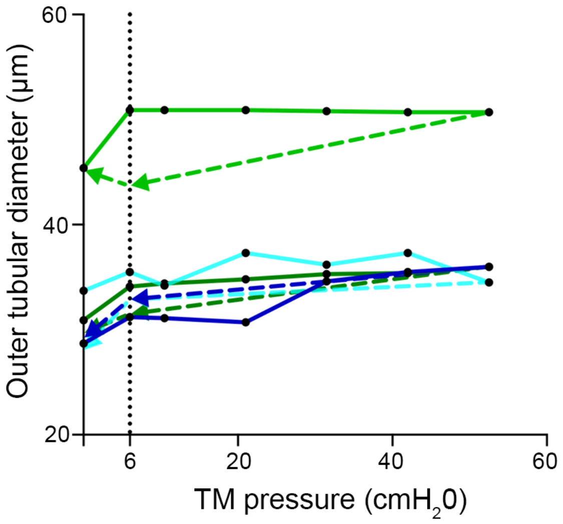


**Extended data figure 5. Control and *Pkd1*-deficient tubules display an elastic behavior.** Once luminal pressure has been increased (solid lines), releasing the pressure (dotted line) reduces tubules diameters. The diameter-transmural (TM) pressure curves of isolated collecting ducts from 8-week-old control (n=2; blue) and *Pkd1*$^{\Delta tub}$ (n=2; green) mice are represented.

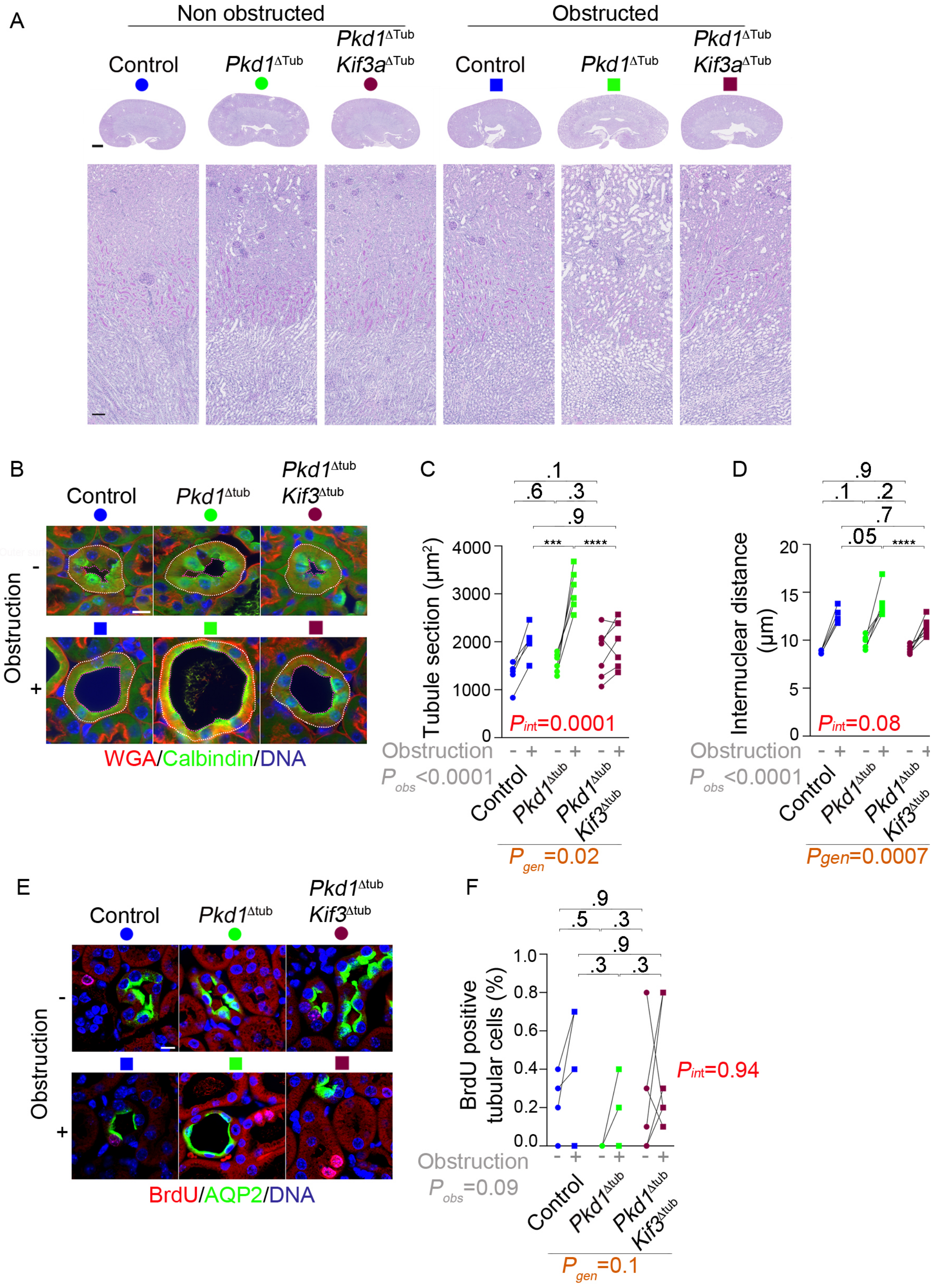


**Extended data figure 6**

**Extended data figure 6. Ureteral obstruction accelerates Polycystin-1 deficient nephron dilation in a cilia-dependent manner.** (**A**) Periodic-acid Schiff's (PAS) staining of obstructed and non-obstructed kidneys from 8-week-old control, *Pkd1*$^{\Delta tub}$ and *Pkd1*$^{\Delta tub}$; *Kif3a*$^{\Delta tub}$ one day after unilateral ureteral obstruction (UUO). Scale bars: 1 mm (up) and 0.1 mm (down). (**B**) Labelling of DNA, wheat germ agglutinin (WGA, which stains the brush border of proximal tubule [PT] and all basement membranes) and calbindin (a distal convoluted tubule [DCT] marker) of obstructed and non-obstructed kidneys from 8-week-old control, *Pkd1*$^{\Delta tub}$ and *Pkd1*$^{\Delta tub}$; *Kif3a*$^{\Delta tub}$ mice one day after unilateral ureteral obstruction (UUO). Scale bar: 10 µm. (**C-D**) Quantification of the mean tubule cross-sectional area (D) and the internuclear distance in DCT (D) of kidneys from the same groups of animals. (**E-F**) labelling of bromodeoxyuridine (BrdU), DNA and aquaporin2 (AQP2; a collecting duct marker; E) and quantification (F) of BrdU staining in collecting ducts of obstructed and non-obstructed kidneys from 8-week-old control, *Pkd1*$^{\Delta tub}$ and *Pkd1*$^{\Delta tub}$; *Kif3a*$^{\Delta tub}$ mice one day after unilateral ureteral obstruction (UUO). Scale bar: 10 µm. Each pair of linked symbols (dot and square) represents the obstructed (square) and non-obstructed (dot) kidneys of an individual female mouse. Two-way ANOVA, P value for obstruction (grey; $P_{obs}$), genotype (brown; $P_{gen}$) and their interaction (red; $P_{int}$), followed by Tukey-Kramer test: *P<0.05, **P<0.01, ***P<0.001, ****P<0.0001 or the indicated P-value.

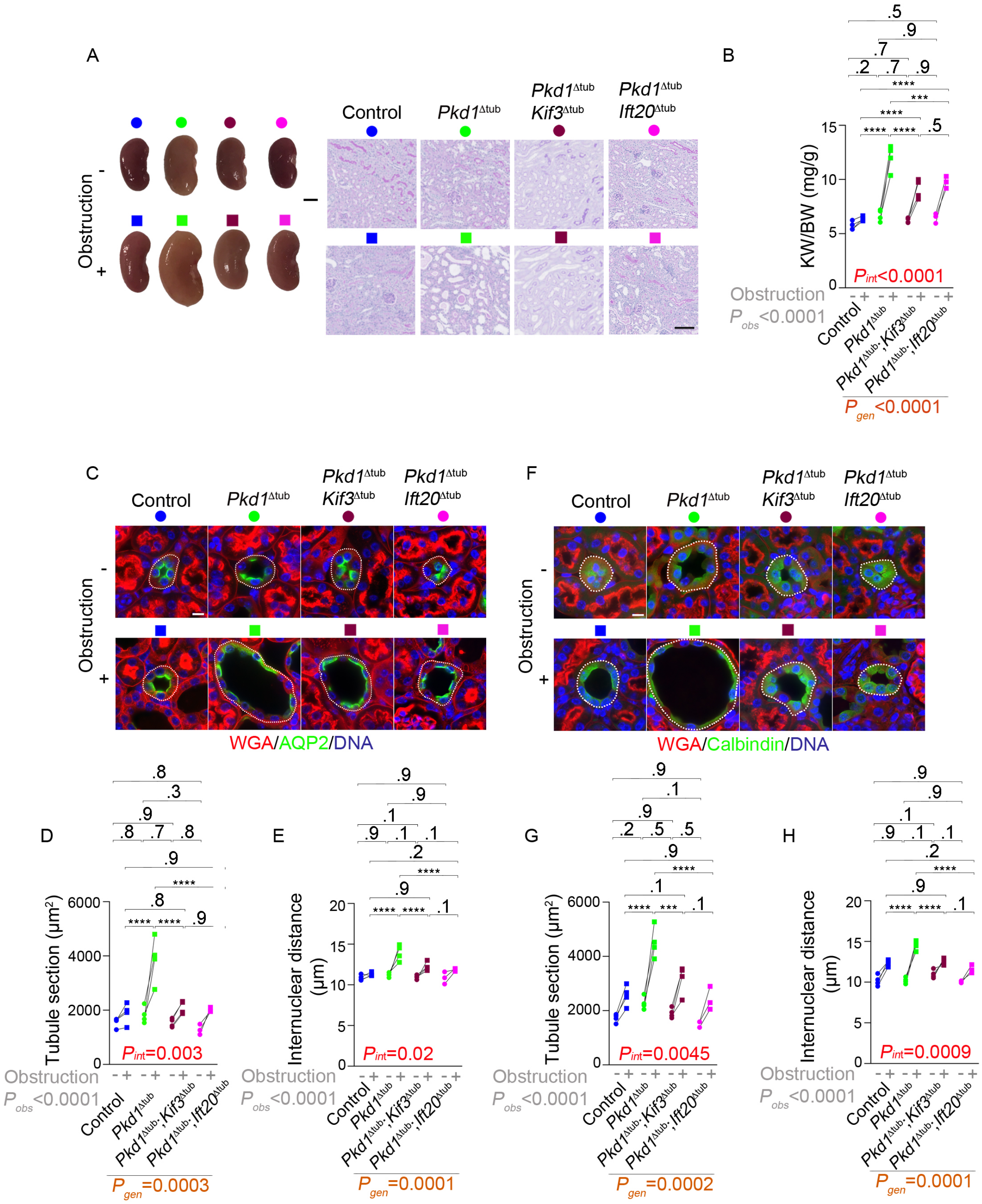


**Extended data figure 7**

**Extended data figure 7. Ureteral obstruction triggers explosive cystogenesis in *Pkd1*$^{\Delta tub}$ distal tubules in a cilia-dependent manner.** (**A-B**) Kidney pictures and Periodic-acid Schiff's staining (A) and quantification (B) of kidney weight to body weight ratio (KW/BW) of obstructed and non-obstructed kidneys from 8 weeks-old control, *Pkd1*$^{\Delta tub}$, *Pkd1*$^{\Delta tub}$; *Kif3a*$^{\Delta tub}$ and *Pkd1*$^{\Delta tub}$; *Ift20*$^{\Delta tub}$ mice four days after unilateral ureteral obstruction (UUO). Scale bars: 1 mm (left) and 0.1 mm (right). Each dot represents one individual female mouse. (**C-H**) labelling of DNA, wheat germ agglutinin (WGA, which stains the brush border of proximal tubule [PT] and all basement membranes), aquaporin 2 (AQP2, a collecting duct [CD] marker; C) or calbindin (a distal convoluted tubule [DCT] marker; F) and quantification of the mean tubule cross-sectional area (D, G) and internuclear distance (E, H) of CD (D-E) and DCT (G, H) in obstructed and non-obstructed kidneys from control, *Pkd1*$^{\Delta tub}$, *Pkd1*$^{\Delta tub}$; *Kif3a*$^{\Delta tub}$ and *Pkd1*$^{\Delta tub}$; *Ift20*$^{\Delta tub}$ mice four days after UUO. Each pair of linked symbols (dot and square) represents the obstructed (square) and non-obstructed (dot) kidneys of an individual female mouse. Scale bars: 10 µm. Two-way ANOVA, P value for obstruction (grey; $P_{obs}$), genotype (brown; $P_{gen}$) and their interaction (red; $P_{int}$), followed by Tukey-Kramer test: *P<0.05, **P<0.01, ***P<0.001, ****P<0.0001 or the indicated P-value.

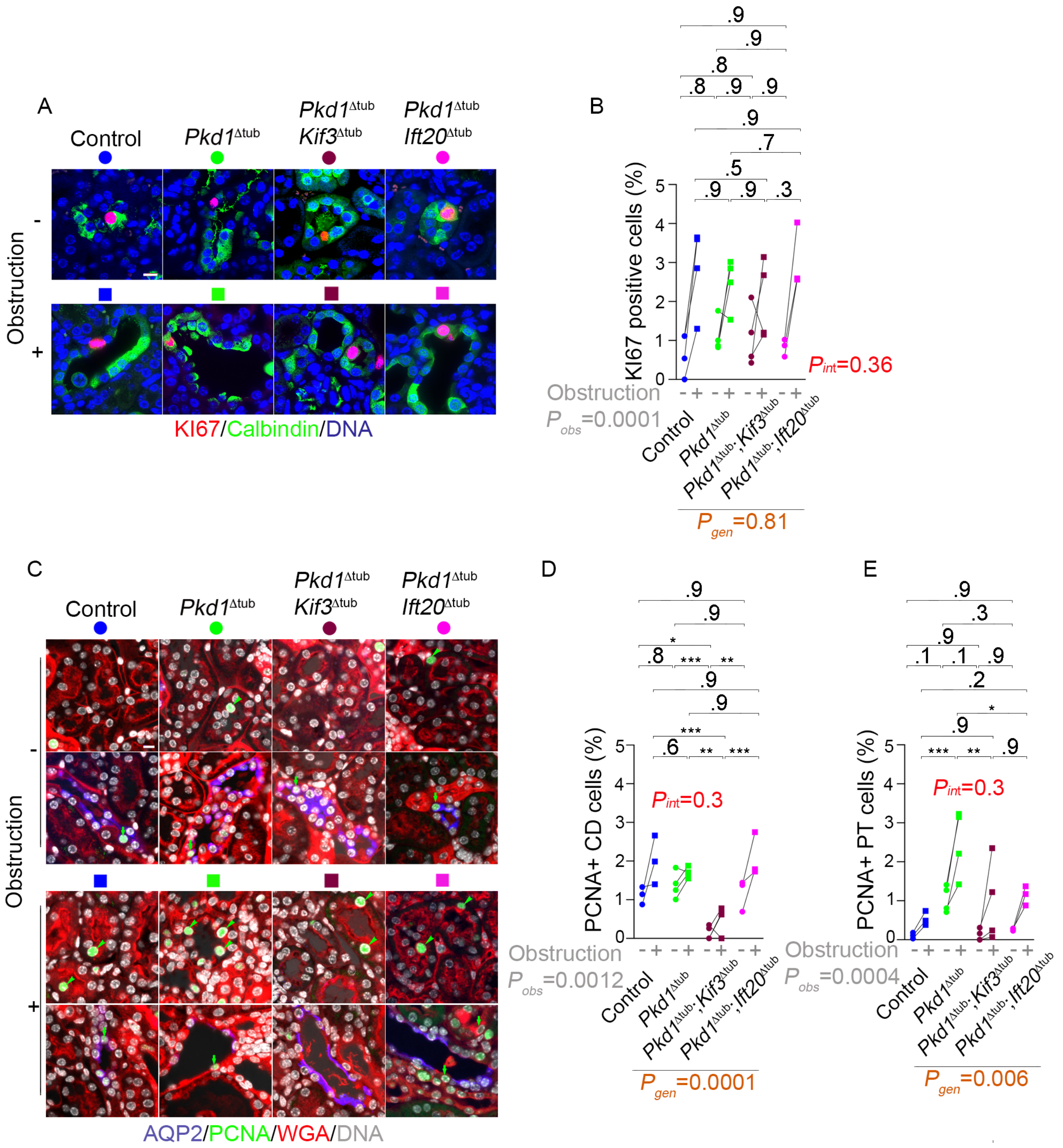
A
Control
Pkd1Δtub
Pkd1Δtub Kif3Δtub
Pkd1Δtub Ift20Δtub
Obstruction
-
+
KI67/Calbindin/DNA
B
KI67 positive cells (%)
Obstruction
Pobs=0.0001
Pint=0.36
Pgen=0.81
Control
Pkd1Δtub
Pkd1Δtub;Kif3Δtub
Pkd1Δtub;Ift20Δtub
C
AQP2/PCNA/WGA/DNA
D
PCNA+ CD cells (%)
Pint=0.3
Pobs=0.0012
Pgen=0.0001
E
PCNA+ PT cells (%)
Pint=0.3
Pobs=0.0004
Pgen=0.006

**Extended data figure 8. Variation in cell proliferation does not explain cilia-dependent *Pkd1*$^{\Delta tub}$ tubule dilation after obstruction**. (**A-B**) Labelling (A) of KI67, DNA, and calbindin (a distal convoluted tubule [DCT] marker) and quantification (B) of KI67 staining in DCT of obstructed and non-obstructed kidneys from control, *Pkd1*$^{\Delta tub}$, *Pkd1*$^{\Delta tub}$; *Kif3a*$^{\Delta tub}$ and *Pkd1*$^{\Delta tub}$; *Ift20*$^{\Delta tub}$ mice, four days after unilateral ureteral obstruction performed at 8 weeks of age. (**C-E**) Representative labelling (C) of proliferation cell nuclear antigen (PCNA), DNA, wheat germ agglutinin (WGA, which stains the brush border of proximal tubule [PT] and all basement membranes) and aquaporin 2 (AQP2, a collecting duct [CD] marker) and quantification of PCNA staining in CD (D; green arrows) and PT (E; green arrowheads) of obstructed and non-obstructed kidneys from the same animals. Each pair of linked symbols (dot and square) represents the obstructed (square) and non-obstructed (dot) kidneys of an individual female mouse. Scale bars: 10 µm. Two-way ANOVA, P value for obstruction (grey; $P_{obs}$), genotype (brown; $P_{gen}$) and their interaction (red; $P_{int}$), followed by Tukey-Kramer test: *P<0.05, **P<0.01, ***P<0.001, ****P<0.0001 or the indicated P-value.
